# ER and SOCE Ca^2+^ signals are not required for directed cell migration in human microglia

**DOI:** 10.1101/2024.01.18.576126

**Authors:** Alberto Granzotto, Amanda McQuade, Jean Paul Chadarevian, Hayk Davtyan, Stefano L. Sensi, Ian Parker, Mathew Blurton-Jones, Ian Smith

## Abstract

The central nervous system (CNS) is constantly surveilled by microglia, highly motile and dynamic cells deputed to act as the first line of immune defense in the brain and spinal cord. Alterations in the homeostasis of the CNS are detected by microglia that respond by migrating toward the affected area. Understanding the mechanisms controlling directed cell migration of microglia is crucial to dissect their responses to neuroinflammation and injury. We used a combination of pharmacological and genetic approaches to explore the involvement of calcium (Ca^2+^) signaling in the directed migration of induced pluripotent stem cell (iPSC)-derived microglia challenged with a purinergic stimulus. This approach mimics cues originating from injury of the CNS. Unexpectedly, simultaneous imaging of microglia migration and intracellular Ca^2+^ changes revealed that this phenomenon does not require Ca^2+^ signals generated from the endoplasmic reticulum (ER) and store-operated Ca^2+^ entry (SOCE) pathways. Instead, we find evidence that human microglial chemotaxis to purinergic signals is mediated by cyclic AMP in a Ca^2+^-independent manner. These results challenge prevailing notions, with important implications in neurological conditions characterized by perturbation in Ca^2+^ homeostasis.

## Introduction

Microglia, the resident immune cells of the central nervous system (CNS), are highly dynamic cells that constantly survey the microenvironment to ensure a rapid response to injuries, infections, or pathological challenges [1,2]. Their distinctive branched morphology is particularly suited for sampling the parenchyma of the brain and the spinal cord for molecular cues suggestive of disruptions in homeostasis of the CNS. Perturbations triggered by an acute injury or chronic neurodegenerative conditions are sensed by microglia that respond by moving toward the damaged site. In this regard, directed migration is essential to position microglia in close proximity to injury or pathologically disrupted areas, where the cells can engage in a range of functions, including phagocytosis, antigen presentation, and cytokine and chemokine release [3].

Purinergic signaling is critical for microglia functioning, and alterations of this specific pathway have been associated with changes in the functional and transcriptomic profiles of microglia in disease-relevant settings [4,5]. Accumulating evidence indicates that the ATP and ADP nucleotides, released from damaged cells, act as key modulators of the purinergic signaling cascade that mediates microglial responses, including process extensions and migration [6–9]. Purinergic receptors, specifically the P2Y_12_ and P2Y_13_ receptor subtypes, are highly expressed in microglia and are activated by ADP [10,11]. These G-protein coupled receptors modulate intracellular signaling events that help orchestrate the transition of microglia towards non-homeostatic states, regulate microglial branching and surveillance, and promote directed migration towards ATP/ADP-mediated chemotactic stimuli [4,6–8,12–14].

Similarly to many other cell types of the immune system, microglia employ calcium ions (Ca^2+^) as a second messenger to signal cues from the extracellular space to the intracellular milieu [1]. Cytosolic Ca^2+^ changes in response to purinergic challenges primarily originate from a signaling cascade that involves the generation of inositol trisphosphate (IP_3_) and the subsequent IP_3_-dependent release of Ca^2+^ from the endoplasmic reticulum (ER). Depletion of Ca^2+^ from the ER activates store-operated Ca^2+^ channels (CRAC), formed by the interaction between STIM1 and ORAI proteins [15]. This store-operated Ca^2+^ entry (SOCE) mechanism contributes to replenishing the lumen of the ER in non-excitable cell types, including microglia [1,15]. The combined activation of these routes of Ca^2+^ signaling results in a biphasic change in calcium (Ca^2+^_i_) levels – an early sharp peak followed by a sustained elevation of Ca^2+^_i_ lasting several seconds [7]. The contribution of Ca^2+^ signals to rodent microglia functioning has been extensively studied in *in vitro* and *in vivo* settings and in the context of pathological conditions [16–19]. However, the impact and precise patterns of Ca^2+^ signals underlying the regulation of directed microglia migration following purinergic receptor activation remain incompletely understood. This issue might also have important pathological implications. Indeed, dysregulation of Ca^2+^ signaling is a pleiotropic mechanism of cellular demise common to acute and chronic neurological conditions, including stroke, Alzheimer’s disease, Parkinson’s disease, and Huntington’s disease [20]. Of note, while defects in neuronal Ca^2+^ signaling are widely recognized as pathogenetic factors in CNS disorders, the impact of these alterations on brain-resident immune cells has received considerably less attention [21].

Primary cell cultures derived from rodent models have been the tools of choice for investigating the pathophysiological features of the cells of the CNS. However, growing transcriptomic and functional evidence indicates that critical species-specific differences exist among the different cell types of the CNS, with variations being prominent in non-neuronal cells like microglia [22,23]. Of note, microglia species-related differences appear to be further exacerbated in pathological settings, likely when the cells are engaged to counteract disease-associated challenges [24,25], thereby limiting the translation of current findings to human-related settings. Novel approaches have been developed to overcome these limitations by generating microglia from human-induced pluripotent stem cells (iPSC), termed iPSC-microglia or iMG [12,26,27]. These cells demonstrate transcriptomic and functional attributes closely akin to those of cultured human microglia [26–28]. Given their utility for examining the impact of disease risk genes on human microglia function and their significant potential in cell-based therapies, they have garnered considerable attention [7,29,30]. However, the mechanisms guiding their directed movement remain largely underexplored. Thus, we leveraged iMG to investigate the role of purinergic signaling-driven Ca^2+^ changes in controlling directed microglia migration.

## Materials and methods

### Generation of iPSCs from human fibroblasts

Human iPSC lines were generated by the University of California, Irvine–Alzheimer’s Disease Research Center (UCI ADRC) Induced Pluripotent Stem Cell Core from subject fibroblasts [31]. Protocols and procedures were approved by the Institutional Review Boards (IRB) and human Stem Cell Research Oversight (hSCRO) committee and informed consent was obtained from all individuals who donated fibroblasts. Non-integrating Sendai virus technology was employed for fibroblast reprogramming. iPSC lines were validated by Array Comparative Genomic Hybridization assessment of karyotype and copy number variation (Cell Line Genetics) and confirmed to be sterile and pluripotent via MycoAlert testing (Lonza), Pluritest Array Analysis, and trilineage in vitro differentiation. iPSCs were cultured on Matrigel (Corning) in antibiotic-free complete mTeSR1 or TeSR-E8 media (StemCell Technologies) in a humidified incubator (5% CO_2_, 37° C).

### CRISPR-editing of iPSCs

The ORAI1 CRISPR-knockout line was generated as previously described [7]. Briefly, 2.5 × 10^5^ iPSCs (UCI ADRC76) digested for 3 min at 37 °C in Accutase and then resuspended in 60 μL nucleofection buffer from the Human Stem Cell Nucleofector™ Kit 2 (Lonza). The suspension was combined 2 μM Electroporation Enhancer (IDTDNA) and 50 μg of RNP complex formed by incubating Alt-R® S.p. HiFi Cas9 Nuclease V3 (IDTDNA) with fused crRNA (5’CGCUGACCACGACUACCCAC):tracrRNA (IDTDNA) duplex for 15 min at 23 °C. The suspension was transferred to the Amaxa Nucleofector cuvette and transfected using program B-016. Cells were plated in TeSR™-E8™ (StemCell Technologies) media with 0.25 μM Thiazovivin and CloneR™ supplement (StemCell Technologies) overnight to recover. Cells were digested the following day with Accutase and single-cell plated to 96-well plates in TeSR™-E8™ media with 0.25 μM Thiazovivin and CloneR™ supplement (StemCell Technologies) for clonal isolation and expansion as previously described [32]. Genomic DNA was extracted using Extracta DNA prep for PCR (Quantabio) from a sample of each clone upon passage and amplified for sequencing using Taq PCR Master Mix (ThermoFisher Scientific) at the cut site using the following primers: ORAI1_F (5’GTAGGGCTTTCTGCCACTCT) and ORAI1_R (5’TATGGCTAACCAGTGAGCGG). PCR product from identified clones was transformed using TOPO™ TA Cloning™ Kit for Subcloning, with One Shot™ TOP10 (ThermoFisher Scientific) for allele-specific sequencing.

The *SALSA6f* line was similarly generated as previously described [7]. Briefly, 2.5 × 10^5^ iPSCs were combined with 1μg of plasmid template and 50 μg of RNP complex targeting the AAVS1 safe harbor locus (crRNA 5’GGGGCCACUAGGGACAGGAU). Clonal isolation and expansion were similarly performed using the following primers to confirm biallelic integration of the *SALSA6f* construct: SV40_F (5’CCACAACTAGAATGCAGTGAA), AAVS1_R (5’GGCTCCAUCGTAAGCAAACC), Puro_R (5’ GTGGGCTTGTACTCGGTCAT), and AAVS1_F (5’ CGGGTCACCTCTCACTCC).

### iMG differentiation

iMG were generated as described in [27]. Briefly, iPSCs were differentiated towards a hematopoietic lineage with the STEMdiff Hematopoiesis kit (StemCell Technologies). After 10-12 days, CD43+ hematopoietic progenitor cells were harvested and moved to a defined, serum-free microglia differentiation medium containing DMEM/F12, 2× insulin-transferrin-selenite, 2× B27, 0.5× N2, 1× Glutamax, 1× non-essential amino acids (NEAA), 400 μM monothioglycerol, and 5 μg/mL human insulin. Cultures were fed every other day with fresh medium supplemented with 100 ng/mL IL-34, 50 ng/mL TGF-β1, and 25 ng/mL M-CSF (Peprotech) for 28 days, eventually frozen in Bambanker freezing medium (2 million iMG cells/vial) and stored in liquid nitrogen. iMG cells were thawed and let recover for one week in complete differentiation medium. In the final 3 days, 100 ng/mL CD200 (Novoprotein) and 100 ng/mL CX3CL1 (Peprotech) were added to further mimic a homeostatic brain environment. Tree independent iPSC lines were used in the study: the ADRC5, the ADRC76, and the Salsa6f lines [7,27].

### Simultaneous recording of intracellular calcium changes and directed cell migration

To monitor iMG migration towards chemoattractants, a micropipette-based assay was implemented as described elsewhere [33], with modifications. In brief, fully differentiated iMG, expressing the genetically encoded ratiometric Ca^2+^ indicator Salsa6f [34], were seeded onto fibronectin-coated glass-bottom petri dishes (Mattek) at a density of 20,000 – 30,000 cells / cm^2^. 18 h after plating, cells were gently washed and bathed in a HEPES-buffered salt solution (HBSS; Sigma-Aldrich) whose composition was (in mM): 135 NaCl, 5.4 KCl, 1.0 MgCl_2_, 10 HEPES, 10 glucose, 2.0 CaCl_2_, and pH 7.4 at 37° C. Ca^2+^-free experiments were performed by omitting Ca^2+^ from the HBSS solution and by supplementing the medium with EGTA (0.3 mM; Sigma-Aldrich). Perturbation of Ca^2+^_i_ levels was performed by loading iMG with the cell-permeable Ca^2+^ chelators BAPTA-AM and EGTA-AM. Briefly, iMG were loaded with either BAPTA-AM (20 μM or 5 μM) or EGTA-AM (20 μM) plus 0.1% Pluronic F-127 for 20 min at 37° C in microglia culture medium and then incubated for further 30 min at room temperature (RT) in HBSS. Control sister cultures were treated with vehicle and followed the same procedure. A thin film of biocompatible silicone oil (Ibidi) was layered on top to prevent medium evaporation. Cells were subsequently transferred to a 37° C heated-stage (Okolab) mounted on an inverted microscope (Nikon Eclipse T*i*, Nikon) equipped with a 10x objective (N.A.: 0.30, Nikon), and a computer-controlled emission filter wheel (Sutter Instruments). To visualize Salsa6f, 488 nm and 560 nm diode lasers (Vortran Laser Technologies) were employed for sequential excitation of GCaMP6f and TdTomato, respectively. Emission of the two fluorescent proteins was collected through a quad band filter and the GCaMP6f signal further filtered through a 524/45 nm band-pass filter. After baseline acquisition (2 to 5 min), a stable chemotactic gradient was generated by pulsing a chemoattractant (ADP 50 µM in the pipette solution, Sigma-Aldrich) from the tip of a pulled, glass pipette placed at the center of the imaging field. The frequency and duration of the chemotactic puffs (0.5 Hz and 20 ms, respectively) were controlled with a Picospritzer II (Parker Instruments) coupled to an external stimulator (Grass Instruments). Stability and consistency of the gradient was monitored throughout the experimental sessions by supplementing the chemotactic solution with Alexa633 (100 nM, ThermoFisher). The dye was excited with a 633 nm diode laser (Vortran Laser Technologies) and emission collected. GCamp6f, TdTomato, and Alexa633 fluorescence images were sequentially acquired with an Orca Flash 4.0LT CMOS camera (Hamamatsu) with a bit depth of 16 bits, using 2 × 2 binning, and cropped for a final field at the specimen of 650 × 650 pixels (one binned pixel = 1.3 µm) at rates of 0.3 frames s^−1^ (Suppl. Movie 1). Image data acquisition and analysis were performed using Nikon Elements NIS software (Nikon).

### IP_3_ uncaging experiments

Uncaging of i-IP_3_ was performed as previously described [7], with modifications. Briefly, iMG were loaded by incubation with the cell-permeable, caged i-IP_3_ analog ci-IP_3_/PM (1 μM, SiChem) plus 0.1% Pluronic F-127 for 20 min at 37° C in microglia culture medium. Cells were gently washed and incubated for an additional 30 min at room temperature (RT) in HBSS medium. During directed cell migration experiments, ci-IP_3_ was uncaged by exposing the imaged cells to two 1-s flashes of ultraviolet (UV) light (350–400 nm) from a xenon arc lamp. UV flash duration was controlled by an electronic shutter (Uniblitz). Effective release of i-IP_3_ was confirmed by increased GCamP6f / TdTomato ratio after the UV flash.

### Calcium imaging experiments

Analysis of intracellular Ca^2+^_i_ changes in iMG expressing the Salsa6f sensor were performed as previously described, with some modifications [7]. After background subtraction, regions of interest (ROI) were drawn for each cell with a semi-automated ROI identification tool. The GCamP6f/TdTomato fluorescence ratio R was calculated for each ROI at each time point. Normalized single-cell ratio values, expressed as ΔR/R (where R_0_ is the mean fluorescence ratio before chemoattractant application and ΔR the fluorescence change (R − R_0_) over time), were used to calculate Ca^2+^ amplitude, a proxy of maximum cation load, and Ca^2+^ integral, an index of overall cation load [7,35]. The analysis of Ca^2+^ changes was limited to the first 100s following the chemoattractant challenge to avoid confounding due to excessive cell branching and need for ROI reshaping during cell movement.

Quantification of Ca^2+^ changes in iMG lacking the Salsa6f construct was performed using a cell-permeable synthetic Ca^2+^ indicator, as previously described [7,36]. Briefly, iMG were loaded by incubation with Cal-590 AM (5 μM, AAT Bioquest) plus 0.1% Pluronic F-127 for 20 min at 37° C in microglia culture medium and then incubated for further 30 min at room temperature (RT) in HBSS. Experiments were performed at RT on the same rig employed for directed cell migration. Images were acquired with a 16-bit Orca Flash 4.0LT CMOS camera (Hamamatsu) with a 2 × 2 binning and cropped to a final field of view of 650 × 650 pixels (one binned pixel = 1.3 µm). Images were acquired at a rate of 1 frame s^−1^. Nikon Elements NIS software (Nikon) was employed for image data acquisition and analysis. Background was subtracted from every frame in the time series, and changes in Cal-590 fluorescence were expressed as ΔF/F_0_, where F_0_ is the resting fluorescence intensity and ΔF is the relative fluorescence change (F – F_0_) over time. Ca^2+^ amplitude and integral were calculated as described above. Store-operated Ca^2+^ entry (SOCE) rate was calculated as ΔF/Δt(s^−1^) over a 10 s time frame after Ca^2+^ supplementation.

### Simultaneous recording of WT and ORAI1 KO iMG cell migration

WT and *ORAI1* KO iMG cells were loaded with two spectrally distinct dyes to monitor the directed migration performances of the two lines within the same dish simultaneously. WT and *ORAI1* KO iMG were collected and loaded, in suspension, with CellTracker Green CMFDA and CellTracker Orange CMTMR (10 μM, Invitrogen) in a culture medium at 37° C for 15 minutes. After two washes in PBS the two cell lines were combined in a 1:1 ratio and plated onto fibronectin-coated glass-bottom petri dishes (Mattek) at a density of 10,000 – 15,000 cells per line / cm^2^. The CellTracker dyes were sequentially excited with 488 nm and 560 nm diode lasers (Vortran Laser Technologies), and emission signals were collected as described above (Suppl. Movie 2). Image sets were acquired every 15 s with the Nikon Elements NIS software (Nikon) and stored for offline analysis.

### Cell tracking and migration analysis

A workflow for the analysis of iMG migration/motility was developed. Starting 1 min after the release of the chemoattractant, 10 min-long time course image series (.nd2 files) were trimmed from the full-length experiment and downsampled to 1-acquisition every 15 s by removing extra images. After background subtraction, time series were analyzed with a custom-made NIS-Elements General Analysis (GA3) routine that included the following steps: segmentation, masking, and cell tracking. The final output was manually curated, and false-positive and false-negative traces were removed. Only cells tracked for at least 20 consecutive acquisitions (5 min) were included in the subsequent analysis. Raw data were exported in the .xlsx format, sorted in MATLAB (MathWorks), and further analyzed with Microsoft Excel (Microsoft).

### Statistical analysis

Microsoft Excel (Microsoft) and OriginPro (OriginLab) were employed for statistical testing and data plotting. Data are represented as mean ± 1 standard error of the mean (s.e.m.). By conventional criteria, differences were considered statistically significant when *p* < 0.05. Details of the number of replicates and the specific statistical test used are provided in the individual figure legends.

## Results

### An ADP gradient promotes directed migration of human microglia via the activation of purinergic signaling

To evaluate the effects of purinergic signaling activation on the migration properties of iMG, we implemented an *in vitro* micropipette-based migration assay ([33] and materials and methods section). The system rapidly generates a stable and consistent chemical gradient, as measured by the signal obtained from the release of the fluorescent dye Alexa 633 (Fig. 1A). To simultaneously monitor the effect of purinergic signaling activation on cell migration and intracellular Ca^2+^ levels, we employed iMG differentiated from an iPSC line engineered to express the genetically-encoded, ratiometric Ca^2+^ indicator Salsa6f [7,34] (Fig. 1B). Two-dimensional representations (’flower plots’) of the cell trajectories during the first 10 minutes after initiating the release of the purinergic agonist ADP (50 µM) at the pipette tip showed a robust migration of iMG towards the center of the chemotactic gradient in a distance- and, thus, concentration-dependent manner (Fig. 1C). To characterize and compare the migration characteristics of iMG, we assessed migration performances by aggregating measurements from cells at 100 µm radial increments along the chemical gradient. Analysis of the mean squared displacement (MSD), a measure of the average distance traveled from their origin by cells over time, showed that ADP stimulation promotes a robust directional migration of iMG located up to 400 µm from the origin of the gradient (Fig. 1D-E and Suppl. Fig. 1A and B). Analysis of the directional migration efficiency, the ratio between the distance traveled towards the center of the gradient and the total distance covered, displayed a similar distance dependence (Fig. 1D-E and Suppl. Fig. 1A and B). Cells located at a greater distance from the pipette (> 400 µm) showed a pattern consistent with Brownian motion (i.e.: a random walk pattern; Fig. 1D-E). Of note, cell speed – defined as the total path length divided by the time elapsed – was not affected by the ADP stimulation and was almost constant regardless of the distance of the cells from the origin of the chemical gradient (Suppl. Fig. 1C). The same set of experiments, performed by omitting ADP from the gradient-generating solution, failed to elicit a directed migratory response, confirming that results are dependent on ADP signaling (Fig. 1I-L and Suppl. Fig. 1F-H).

**Figure 1.**
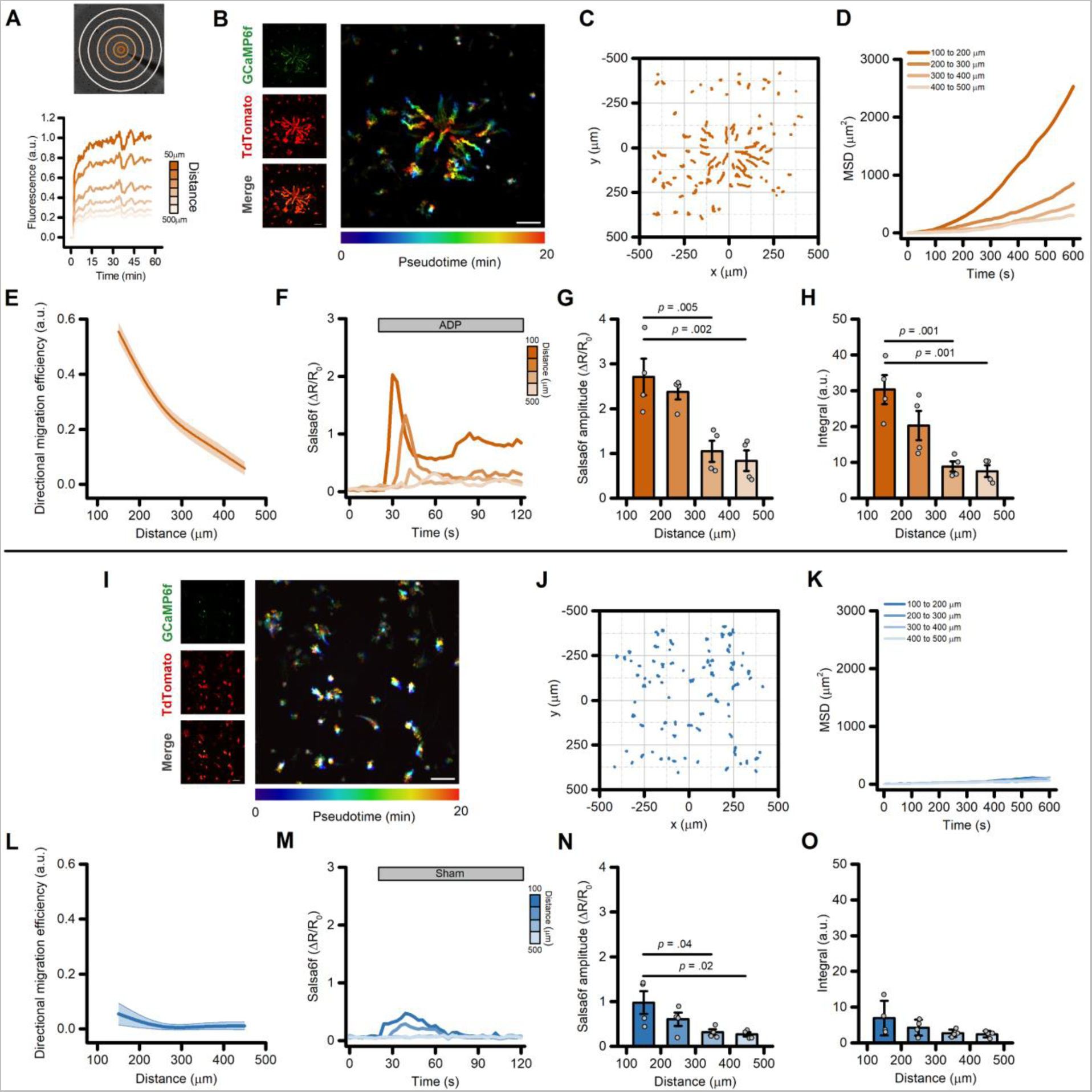
Directed migration of iMG cells following the generation of an ADP gradient. (A) The panel illustrates the experimental paradigm employed to generate and monitor the chemotactic gradient. The *upper panel* shows a brightfield image of iMG cells, overlaid with concentric ROIs used to monitor the stability and consistency of the chemotactic gradient. The *lower panel* shows the time course of measurements of Alexa633 dye fluorescence intensities at increasing distances from the tip of the pipette. Fluorescence data were normalized to the average fluorescence intensity as measured within 50 μm from the tip of the pipette 20-30 min after release of the Alexa633 dye. (B) Maximum intensity projection photomicrographs of Salsa6f-expressing iMG cells challenged with an ADP (50 µM) gradient for up to 20-min. Left panels show, from top to bottom, the GCaMP6f signal, the TdTomato signal, and the corresponding merged image. Right panel, time-course pseudocolored image with warmer colors indicating increased elapsed time. (C) ‘Flower plot’ depicts 10-min migration tracks as analyzed from the iMG cells in B, normalized to the pipette tip coordinates. (D) Mean squared displacement plot of iMG cells exposed to an ADP gradient at 100 µm radial increments (100 to 200 µm: 35 cells; 200 to 300 µm: 58 cells; 300 to 400 µm: 74 cells; 400 to 500 µm: 37 cells). (E) The plot depicts directed migration efficiency towards an ADP gradient averaged at 100 µm radial increments (n = 4 independent experiments). (F) Time course of ADP-dependent Ca^2+^_i_ rises as assessed with the Salsa6f sensor. Traces represent average responses at 100 µm radial increments following ADP release from the pipette (n = 4 independent experiments). (G) Bar graphs depict Ca^2^ ^+^_i_ amplitude values expressed as (ΔR/R_0_). (H) Bar graphs depict Ca^2^ ^+^_i_ integrals, measured as the area under each curve. (I) The images illustrate maximum intensity projection photomicrographs of iMG cells expressing the Salsa6f Ca^2+^ sensor exposed to a sham stimulation. Left panels show, from top to bottom, the GCaMP6f signal, the TdTomato signal, and the corresponding merged image. Right panel shows pseudocolored image across time. Note the lack of migration towards the center of the field of view. (J) ‘Flower plot’ depicts 10-min migration tracks as analyzed from the iMG cells in I, normalized to the pipette tip coordinates. (K) Mean squared displacement plot of iMG cells exposed to sham treatment at 100 µm radial increments (100 to 200 µm: 40 cells; 200 to 300 µm: 63 cells; 300 to 400 µm: 91 cells; 400 to 500 µm: 46 cells). (L) The plot depicts the directed migration efficiency of sham-treated iMG cells averaged at 100 µm radial increments (n = 4 independent experiments). (M) Time course of Ca^2+^_i_ rises as assessed with the Salsa6f sensor and evoked by the sham treatment. Traces represent average responses at 100 µm radial increments following vehicle release from the pipette (n = 4 independent experiments). (N) Bar graphs depict Ca^2^ ^+^_i_ amplitude values expressed as (ΔR/R_0_). (O) Bar graphs depict Ca^2^ ^+^_i_ integrals, measured as the area under each curve. In E and L, a b-spline function was applied for curve smoothing. In G, H, N, and O the comparison of mean values was assessed by one-way ANOVA followed by Tukey’s post-hoc test. Scale bars 100 μm.

Intracellular Ca^2+^ transients displayed a distance-, dose-dependent effect (Fig. 1F), with Salsa6f iMG cells closer to the origin of the ADP gradient showing larger Ca^2+^ amplitudes and Ca^2+^ integrals (Fig. 1G and H, respectively). A modest but significant distance-dependent change was observed in Ca^2+^_i_ levels a few seconds after the beginning of the sham stimulation (Fig. 1N-O), an effect likely due to shear stress forces to which iMG are sensitive [7]. Similar ADP-evoked migration patterns were observed in iMG cells differentiated from a different iPSC line (UCI ADRC5), suggesting the lack of any genetic background- or sex-dependent effect (Suppl. Fig. 1I-L).

We have previously shown that, in iMG cells, ADP primarily signals through the activation of P2Y_12_ and P2Y_13_ purinergic receptors [7,26]. To test the contribution of these two receptors to ADP-evoked directed cell migration, we exposed iMG cells to a combination of P2Y_12_ and P2Y_13_ receptor antagonists (PSB 0739 and MRS 2211, respectively; 10 µM each). This pharmacological approach completely abolished ADP-induced directed cell migration when compared to vehicle-treated sister cultures (Fig. 2A-D and Suppl Fig. 2A). In parallel, the two compounds also abrogated ADP-evoked Ca^2+^_i_ rises (Fig. 2E-H). Of note, inhibition of P2Y_12_ and P2Y_13_ receptors significantly reduced the speed of iMG cells independently of their position along the gradient (Suppl. Fig. 2B), suggesting that tonic activation of purinergic signaling modulates baseline microglia motility.

**Figure 2.**
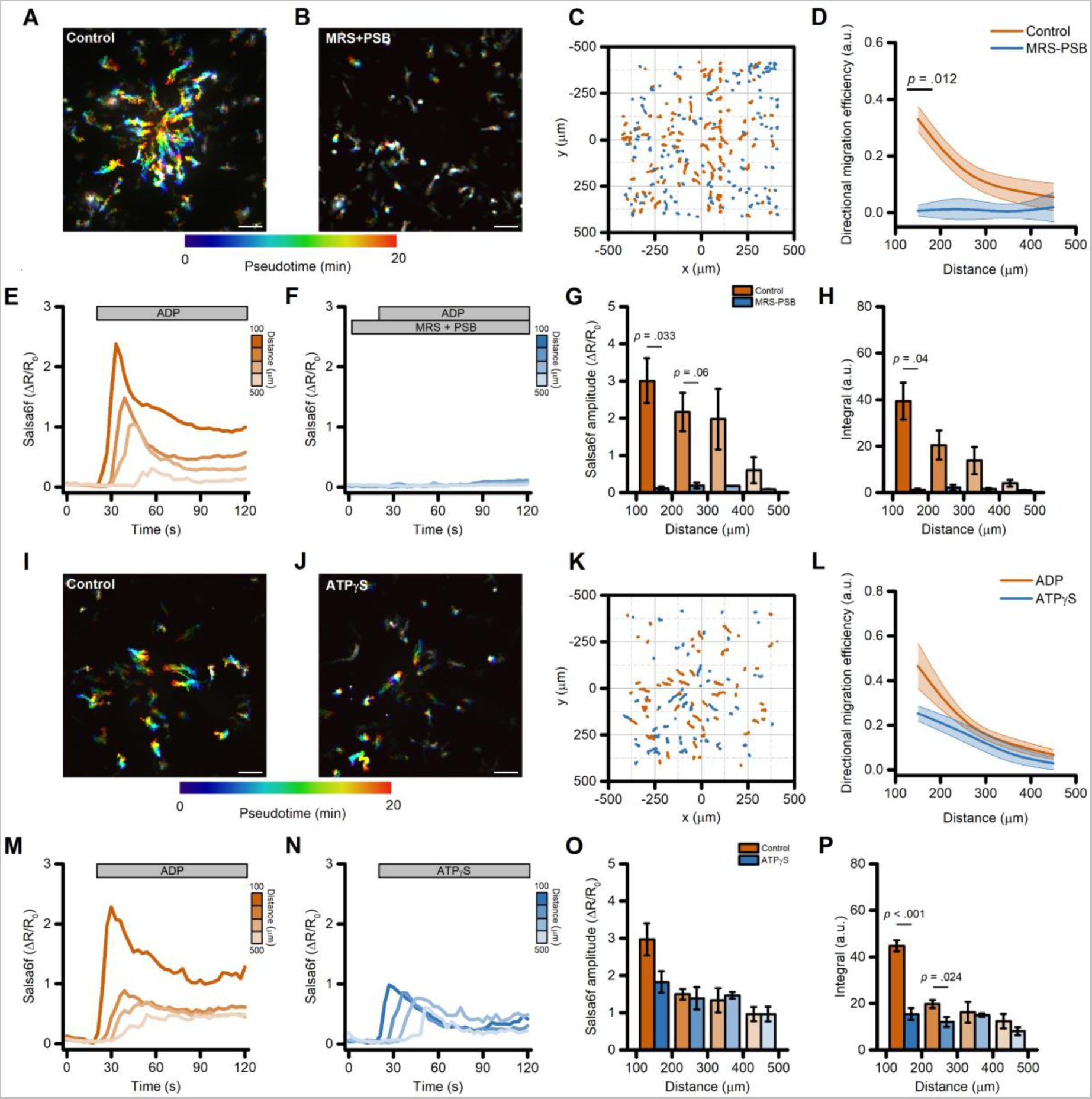
Directed migration of iMG cells is driven by activation of purinergic signaling. (A-B) Pseudocolored maximum intensity projection photomicrographs of iMG cells expressing the Salsa6f sensor and challenged with an ADP gradient in the presence (B) or absence (A) of the P2Y_12_ and P2Y_13_ receptor antagonists PSB 0739 and MRS 2211, respectively. (C) ‘Flower plot’ depicts 10-min migration tracks as analyzed from the iMG cells in A and B; traces from the two populations were normalized to the pipette tip coordinates and overlaid. (D) The plot depicts directed migration efficiency towards an ADP gradient of control- and PSB 0739 + MRS 2211-treated iMG cells averaged at 100 µm radial increments (from n = 3 control and n = 2 PSB 0739 + MRS 2211 independent experiments). (E-F) Time course of ADP-dependent Ca^2+^_i_ rises as assessed with the Salsa6f sensor. Traces represent average responses at 100 µm radial increments following ADP release from the pipette in the presence (F) or absence (E) of PSB 0739 + MRS 2211 (from n = 3 controls and n = 2 PSB 0739 + MRS 2211 independent experiments). (G) Bar graphs depict Ca^2^ ^+^_i_ amplitude values expressed as ΔR/R_0_. (H) Bar graphs depict Ca^2^ ^+^_i_ integrals, measured as the area under each curve obtained in the two populations. (I-J) Pseudocolored maximum intensity projection photomicrographs of iMG cells expressing the Salsa6f sensor and challenged with an ADP (I) or an ATPγS (J) gradient. (K) ‘Flower plot’ depicts 10-min migration tracks as analyzed from the iMG cells in I and J; traces from the two populations were normalized to the pipette tip coordinates and overlaid. (L) Plot depicts directed migration efficiency towards the ADP or the ATPγS gradient of iMG cells averaged at 100 µm radial increments (from n = 5 ADP and n = 2 ATPγS independent experiments). (M-N) Time course of ADP- and ATPγS-dependent Ca^2+^_i_ rises as assessed with the Salsa6f sensor. Traces represent average responses at 100 µm radial increments following ADP (M) or ATPγS (N) release from the pipette (from n = 5 ADP and n = 4 ATPγS independent experiments). (G) Bar graphs depict Ca^2^ ^+^_i_ amplitude values expressed as ΔR/R_0_. (H) Bar graphs depict Ca^2^ ^+^_i_ integrals, measured as the area under each curve obtained in the two populations. In D and L, a b-spline function was applied for curve smoothing. The comparison of mean values was assessed by a two-tailed unpaired Student’s t-test. Scale bars 100 μm.

Directed cell migration of iMG also occurred in response to pipette delivery of similar concentrations of another purinergic agonist ATPγS (50 µM), a non-hydrolyzable ATP analog. Whereas ATP can be readily hydrolyzed to ADP by microglial-expressed ENTPD1/CD39, ATPγS is resistant to this process and thus microglial responses to ATPγS are likely mediated via P2X4 and P2X7 receptors, not P2RY12/13. In comparison to ADP, ATPγS showed a modest, non-significant reduction in migration efficiency (Fig. 2I-L) and a decreased Ca^2+^ response driven primarily by short-lasting Ca^2+^ elevations (Fig. 2M-P) [7]. Other relevant parameters related to cell motility were unaltered when compared to the ADP challenge (Suppl. Fig. 2C-D).

Taken together, these results indicate that the activation of purinergic signaling via P2Y_12_ and P2Y_13_ receptors promotes the robust activation of mechanisms of directional migration in human microglia.

### Perturbation of the extracellular and intracellular Ca^2+^ milieu does not affect directed migration of human microglia

Considering the central role of Ca^2+^ signaling in modulating cell migration, we investigated the contribution of ADP-evoked Ca^2+^_i_ changes in iMG cells during directed cell migration. To test this, we modified the extracellular and intracellular Ca^2+^ milieu and monitored migration and Ca^2+^_i_ dynamics changes in iMG cells exposed to the ADP gradient.

First, we bathed iMG cells in a Ca^2+^-free medium supplemented with the cell impermeable Ca^2+^ chelator EGTA (300 µM) and compared their behavior with sister cultures bathed in a Ca^2+^-containing solution (2 mM). This maneuver resulted in a modest but significant reduction of directed migration efficiency on the cells closer to the gradient’s center (Fig. 3A-D). Similar alterations were observed for other motility parameters, like line speed to the tip (the mean velocity towards the gradient’s center), and straightness, a proxy of the ability of cells to move in a straight line (Suppl. Fig. 3A-B). Speed was unaffected (Suppl. Fig. 3C). Analysis of the Ca^2+^_i_ changes following the ADP challenge showed that initial Ca^2+^ peak amplitude was not significantly different in iMG cells bathed in Ca^2+^-containing or Ca^2+^-free medium (Fig. 3E-G and Suppl. Fig. 3F). On the contrary, the sustained Ca^2+^ phase observed in the presence of external Ca^2+^ was completed abrogated when Ca^2+^ was removed from the solution (Fig. 3E-F and H). These observations are in line with our previous findings [7] and strongly support the idea that the initial Ca^2+^ peak is driven primarily by Ca^2+^ released from intracellular stores, whereas the sustained elevations are supported by Ca^2+^ influx, likely through Ca^2+^ release-activated (CRAC) channels.

**Figure 3.**
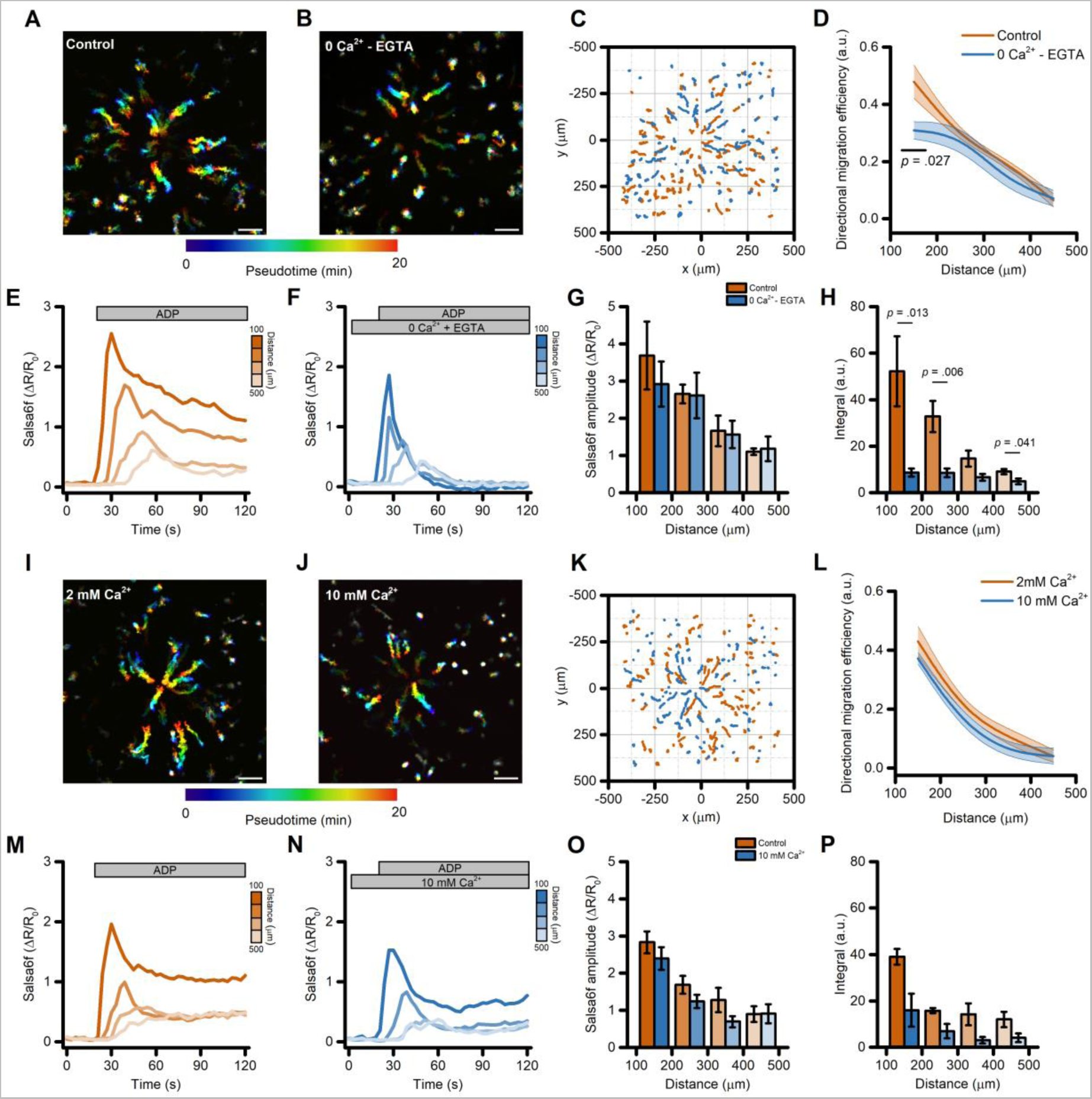
Directed migration of iMG cells is modestly affected by perturbation of the extracellular Ca^2+^ milieu. (A-B) Pseudocolored maximum intensity projection photomicrographs of iMG cells expressing the Salsa6f sensor and bathed in a control (2mM Ca^2+^; A) or a Ca^2+^-free (0 Ca^2+^ + 300 µM EGTA; B) medium and challenged with an ADP gradient. (C) ‘Flower plot’ depicts 10-min migration tracks as analyzed from the iMG cells in A and B; traces from the two populations were normalized to the pipette tip coordinates and overlaid. (D) The plot depicts directed migration efficiency towards an ADP gradient of iMG cells bathed in a control or Ca^2+^-free medium and averaged at 100 µm radial increments (from n = 4 controls and n = 5 Ca^2+^-free independent experiments). (E-F) Time course of ADP-dependent Ca^2+^_i_ rises as assessed with the Salsa6f sensor. Traces represent average responses at 100 µm radial increments following ADP release from the pipette in control (E) or Ca^2+^-free (F) medium (from n = 4 controls and n = 5 Ca^2+^-free independent experiments). (G) Bar graphs depict Ca^2^ ^+^_i_ amplitude values expressed as ΔR/R_0_. (H) Bar graphs depict Ca^2^ ^+^_i_ integrals, measured as the area under each curve obtained in the two populations. (I-J) Pseudocolored maximum intensity projection photomicrographs of iMG cells expressing the Salsa6f sensor and bathed in a control (2mM Ca^2+^; I) or a 10 mM Ca^2+^-containing medium (J) and challenged with an ADP gradient. (K) ‘Flower plot’ depicts 10-min migration tracks as analyzed from the iMG cells in I and J; traces from the two populations were normalized to the pipette tip coordinates and overlaid. (L) The plot depicts directed migration efficiency towards an ADP gradient of iMG cells bathed in a control or 10 mM Ca^2+^-containing medium and averaged at 100 µm radial increments (from n = 5 controls and n = 5 10 mM Ca^2+^ independent experiments). (M-N) Time course of ADP-dependent Ca^2+^_i_ rises as assessed with the Salsa6f sensor. Traces represent average responses at 100 µm radial increments following ADP release from the pipette in control (M) or 10 mM Ca^2+^-containing (N) medium (from n = 5 controls and n = 5 10 mM Ca^2+^ independent experiments). (O) Bar graphs depict Ca^2^ ^+^_i_ amplitude values expressed as ΔR/R_0_. (P) Bar graphs depict Ca^2^ ^+^_i_ integrals, measured as the area under each curve obtained in the two populations. In D and L, a b-spline function was applied for curve smoothing. The comparison of mean values was assessed by a two-tailed unpaired Student’s t-test. Scale bars 100 μm.

To further explore the role of Ca^2+^_i_ signaling during ADP-driven migration, we exposed iMG cells to a medium containing supraphysiological Ca^2+^ concentration (10 mM) and compared their behavior with that in the control (2 mM Ca^2+^) medium. This approach did not affect directed cell migration (Fig. 3I-L) nor any other motility parameters (Suppl. Fig. 3D-E). No significant differences were also observed when analyzing the ADP-evoked Ca^2+^_i_ changes, both in terms of peak amplitude and overall cation load (Fig. 3M-P and Suppl. Fig. 3G), suggesting that iMG cells are well equipped for fast and efficient control of Ca^2+^_i_ homeostasis.

Bulk removal of extracellular Ca^2+^ could have affected cell motility through mechanisms that act independently of changes in Ca^2+^_i_, such as the disruption of integrin-dependent, Ca^2+^-mediated cell anchoring to the extracellular matrix. To circumvent this limitation, we next evaluated the effect of intracellularly-trapped Ca^2+^ chelators on ADP-evoked directed cell migration. No significant changes in migration and motility performances were observed in iMG cells loaded with the fast, high affinity Ca^2+^ chelator BAPTA following incubation with BAPTA-AM (20 µM) [37] when compared to vehicle-treated cultures (Fig. 4A-D and Suppl. Fig. 4A-B). Conversely, ADP-evoked Ca^2+^_i_ rises were significantly delayed, and Ca^2+^_i_ buildup was markedly reduced in BAPTA-loaded cells (Fig. 4E-H and Suppl. Fig. 4C-D). However, it was noticed that at this loading concentration BAPTA had some effects on iMG morphology by the end of the loading procedure (Suppl Fig. 4E). To minimize side effects due to excessive BAPTA accumulation and/or compartmentalization we performed the same set of experiments by reducing the BAPTA-AM loading concentration to 5 µM. At this concentration, BAPTA delayed and reduced the ADP-driven Ca^2+^_i_ changes (Fig. 4M-P and Suppl. Fig. 4H-I), but failed to alter migration (Fig. 4I-L and Suppl. Fig. 4F-G) as compared to vehicle-treated cells.

**Figure 4.**
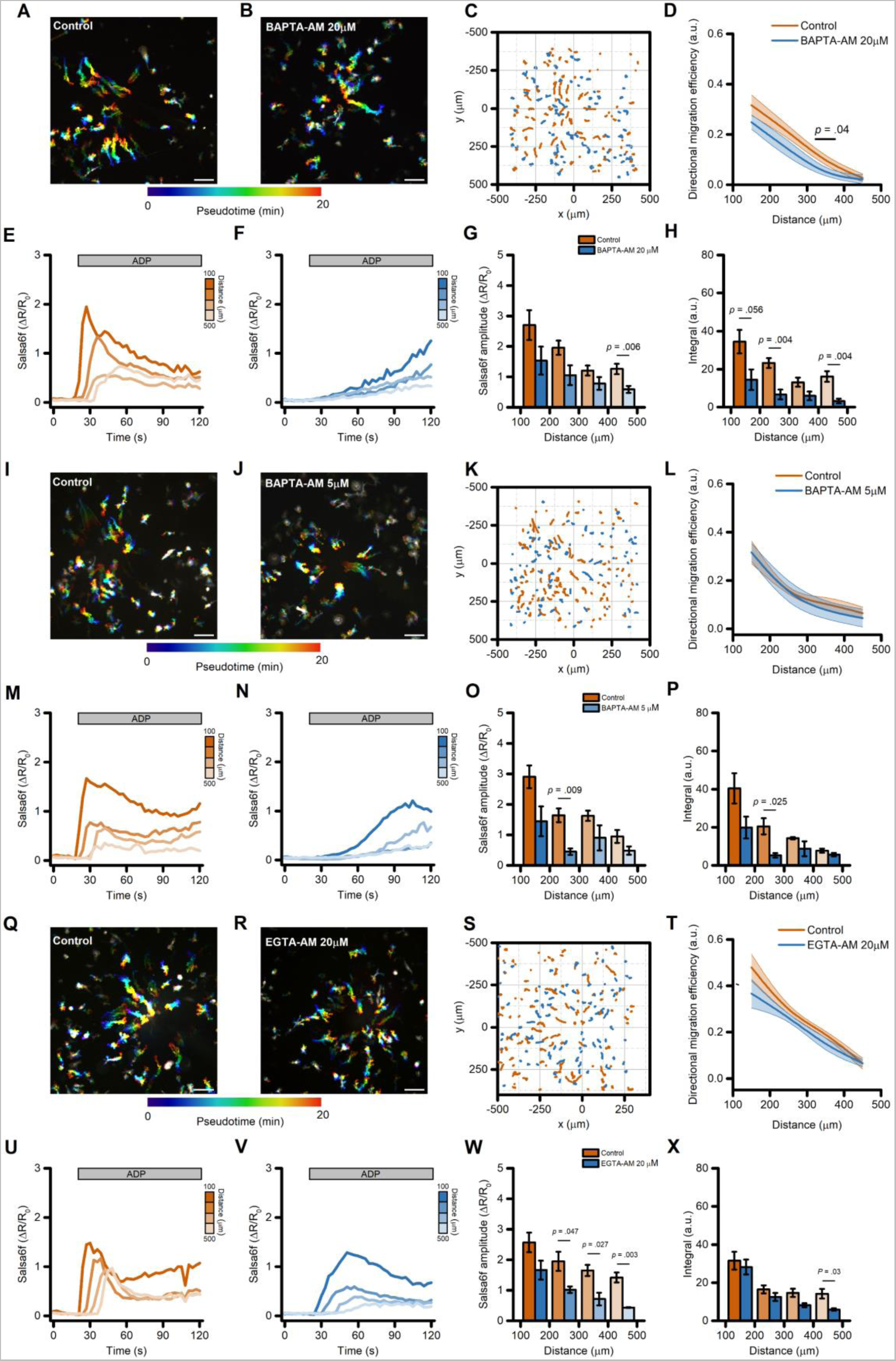
Directed migration of iMG cells is modestly affected by intracellular Ca^2+^ chelators. (A-B) Pseudocolored maximum intensity projection photomicrographs of control (A) and BAPTA-loaded; B) iMG cells expressing the Salsa6f sensor challenged with an ADP gradient. (C) ‘Flower plot’ depicts 10-min migration tracks as analyzed from the iMG cells in A and B; traces from the two populations were normalized to the pipette tip coordinates and overlaid. (D) The plot depicts directed migration efficiency towards an ADP gradient of control and 20 µM BAPTA-loaded iMG cells averaged at 100 µm radial increments (from n = 5 controls and n = 7 BAPTA independent experiments). (E-F) Time course of ADP-dependent Ca^2+^_i_ rises as assessed with the Salsa6f sensor. Traces represent average responses at 100 µm radial increments following ADP release from the pipette in control (E) or BAPTA-loaded (F) iMG cells (from n = 5 controls and n = 7 BAPTA independent experiments). (G) Bar graphs depict Ca^2^ ^+^_i_ amplitude values expressed as ΔR/R_0_. (H) Bar graphs depict Ca^2^ ^+^_i_ integrals, measured as the area under each curve obtained in the two populations. (I-J) Pseudocolored maximum intensity projection photomicrographs of control (I) and BAPTA-loaded (5 µM; J) iMG cells expressing the Salsa6f sensor challenged with an ADP gradient. (K) ‘Flower plot’ depicts 10-min migration tracks as analyzed from the iMG cells in I and J; traces from the two populations were normalized to the pipette tip coordinates and overlaid. (L) The plot depicts directed migration efficiency towards an ADP gradient of control and 5 µM BAPTA-loaded iMG cells averaged at 100 µm radial increments (from n = 3 controls and n = 3 BAPTA independent experiments). (M-N) Time course of ADP-dependent Ca^2+^_i_ rises as assessed with the Salsa6f sensor. Traces represent average responses at 100 µm radial increments following ADP release from the pipette in control (M) or 5 µM BAPTA-loaded (N) iMG cells (from n = 3 controls and n = 3 BAPTA independent experiments). (O) Bar graphs depict Ca^2^ ^+^_i_ amplitude values expressed as ΔR/R_0_. (P) Bar graphs depict Ca^2^ ^+^_i_ integrals, measured as the area under each curve obtained in the two populations. (Q-R) Pseudocolored maximum intensity projection photomicrographs of control (Q) and EGTA-loaded (20 µM; R) iMG cells expressing the Salsa6f sensor challenged with an ADP gradient. (S) ‘Flower plot’ depicts 10-min migration tracks as analyzed from the iMG cells in Q and R; traces from the two populations were normalized to the pipette tip coordinates and overlaid. (T) The plot depicts directed migration efficiency towards an ADP gradient of control and 5 µM EGTA-loaded iMG cells averaged at 100 µm radial increments (from n = 4 controls and n = 4 EGTA independent experiments). (U-V) Time course of ADP-dependent Ca^2+^_i_ rises as assessed with the Salsa6f sensor. Traces represent average responses at 100 µm radial increments following ADP release from the pipette in control (U) or 20 µM EGTA-loaded (V) iMG cells (from n = 4 controls and n = 4 EGTA independent experiments). (W) Bar graphs depict Ca^2^ ^+^_i_ amplitude values expressed as ΔR/R_0_. (X) Bar graphs depict Ca^2^ ^+^_i_ integrals, measured as the area under each curve obtained in the two populations. In D, L, and T a b-spline function was applied for curve smoothing. The comparison of mean values was assessed by a two-tailed unpaired Student’s t-test. Scale bars 100 μm.

Ca^2+^ buffers with different biophysical properties have distinct effects on Ca^2+^_i_ dynamics [37] and could elicit divergent biological responses. We therefore loaded iMG cells with EGTA-AM (20 µM), a Ca^2+^ chelator that has slower buffering kinetics but similar affinity to BAPTA [37,38]. As expected, the ADP-evoked Ca^2+^_i_ dynamics were markedly different between EGTA- *vs.* BAPTA-loaded iMG cells (Fig. 4U-X *vs.* 4E-H). In EGTA-loaded cells exposed to the ADP gradient, Ca^2+^_i_ peak amplitude was significantly reduced when compared to vehicle-treated sister cultures (Fig. 4U-W). Conversely, cumulative Ca^2+^_i_ load and rise time were unaffected (Fig. 4X and Suppl. Fig.4M-N). A comparison of migration performances showed that EGTA loading did not influence the ability of iMG cells to move efficiently along the gradient (Fig. 4T), but significantly affected cell speed (Suppl. Fig. 4J-L).

Altogether, these results show that alterations of Ca^2+^_i_ dynamics have a modest yet significant effect on ADP-evoked directional migration in human microglia.

Altogether, these results show that alterations of Ca^2+^_i_ dynamics modestly impact ADP-induced directional migration in human microglia. Notably, this effect is significant only under reduced extracellular Ca^2+^ conditions, with BAPTA and EGTA experiments suggesting that the impact of external Ca^2+^ may be distinct from its effect on cytosolic Ca^2+^ levels.

### ADP-evoked store-operated Ca2+ entry (SOCE) is not required for directed migration of human microglia

The use of intracellular Ca^2+^ chelators produced only modest changes in Ca^2+^ signals. To overcome this limitation and probe the specific contribution of different Ca^2+^ sources on ADP-evoked directed cell migration, we examined the role of SOCE by using CRISPR gene editing to delete *ORAI1*. In *ORAI1* KO iMG cells loaded with the high-affinity Ca^2+^ indicator Cal-590, SOCE was entirely abrogated upon Ca^2+^ supplementation after depletion of the ER with the SERCA pump inhibitor thapsigargin (1 µM) as compared to wild-type (WT) sister cultures (Fig. 5A-D). Similarly, the deletion of *ORAI1* abolished the SOCE-mediated Ca^2+^ influx triggered by exposure to ADP (Fig. 5E-H). To evaluate the effect of this maneuver on directed migration, WT and *ORAI1* KO cells were labeled independently with two spectrally distinct dyes (CellTracker Green and Orange, respectively) and then combined in the same dish in a 1:1 ratio (Fig. 5I). No significant differences between these cells were observed in ADP-evoked directed cell migration performances (Fig. 5I-J) nor in other motility parameters (Suppl. Fig. 5A-B). Similar results were obtained when WT and *ORAI1* KO cells were bathed in a Ca^2+^-free medium during the migration experiment (Suppl. Fig. 5C-F).

**Figure 5.**
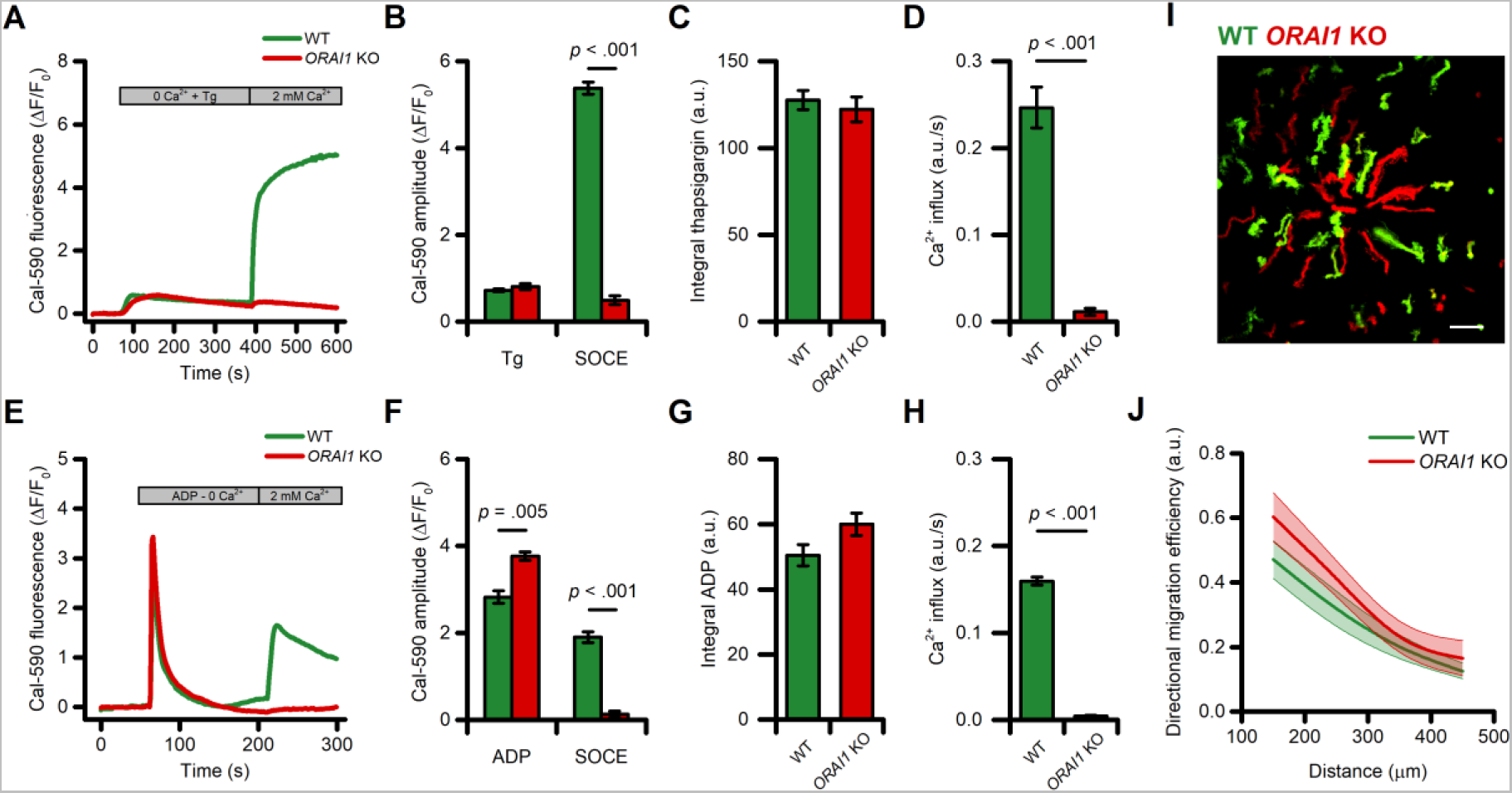
SOCE is not required for directed migration of iMG cells. (A) Time course of SOCE evoked by exposure to thapsigargin (Tg; 1 μM) in WT and *ORAI1* KO cells as assessed with Cal-590 (n = 3 WT and n = 4 *ORAI1* KO independent experiments). (B) Bar graphs depict ER store release (Tg) and SOCE-mediated Ca^2+^ changes (SOCE) quantified as peak amplitude. (C-D) Bar graphs depict SOCE-mediated Ca^2+^_i_ changes quantified as area under the curve (C) and rate of SOCE (D). (E) Time course of SOCE evoked by exposure to ADP (10 μM) in WT and *ORAI1* KO cells as assessed with Cal-590 (n = 3 WT and n = 4 *ORAI1* KO independent experiments). (F) Bar graphs depict Ca^2+^ release from intracellular stores following ADP exposure (ADP) and SOCE-mediated Ca^2+^ changes (SOCE) quantified as peak amplitude. (C-D) Bar graphs depict ADP-mediated Ca^2+^_i_ changes quantified as area under the curve (G) and rate of SOCE (H). (I) Maximum intensity projection photomicrographs of WT (Green) and *ORAI1* KO (Red) iMG cells challenged with an ADP gradient. (J) The plot depicts directed migration efficiency towards an ADP gradient of WT and *ORAI1* KO cells averaged at 100 µm radial increments (from n = 6 independent experiments). In J a b-spline function was applied for curve smoothing. The comparison of mean values was assessed by a two-tailed unpaired Student’s t-test. Scale bar 100 μm.

These findings demonstrate that ORAI1 is necessary for SOCE-driven Ca^2+^ influx in iMG, but the process is dispensable for ADP-evoked directed migration.

### ADP-evoked Ca^2+^ release from intracellular stores is not required for directed migration of human microglia

Experiments performed in Ca^2+^-free medium and in *ORAI1* KO cells showed that the ADP-evoked sustained Ca^2+^ rises are not required for directed migration but do not provide information on the contribution of the early, transient Ca^2+^_i_ elevations originating from the ER [7]. To address this issue, we performed pharmacological manipulations to disrupt Ca^2+^ release from intracellular stores. First, we compared ADP-driven iMG migration before and after uncaging the IP_3_ analog, ci-IP_3_. This approach transiently increased Ca^2+^_i_ levels (Fig. 6A) but failed to affect directed migration along the ADP gradient (Fig. 6B-D). Similarly, additional motility parameters were unaffected (Suppl. Fig. 6A-B). We next evaluated the ADP-evoked Ca^2+^_i_ and migration features of iMG cells exposed to the SERCA pump inhibitor thapsigargin (1 µM, pretreated for 5 min in Ca^2+^-free medium) to fully deplete ER Ca^2+^ content. This maneuver abrogated the ADP-evoked Ca^2+^ release in all cells located along the gradient (Fig. 6J-L), whereas Ca^2+^_i_ changes in vehicle-treated iMG were unaffected (Fig. 6I-L). Notably, thapsigargin administration did not alter directed migration efficiency of iMG along the ADP gradient or any other motility parameters (Fig. 6E-H and Suppl. Fig. 6C-D). Similar results were obtained when iMG cells were pretreated with cyclopiazonic acid (CPA), another SERCA pump inhibitor (Fig. 6M-T and Suppl. Fig. 6E-F).

**Figure 6.**
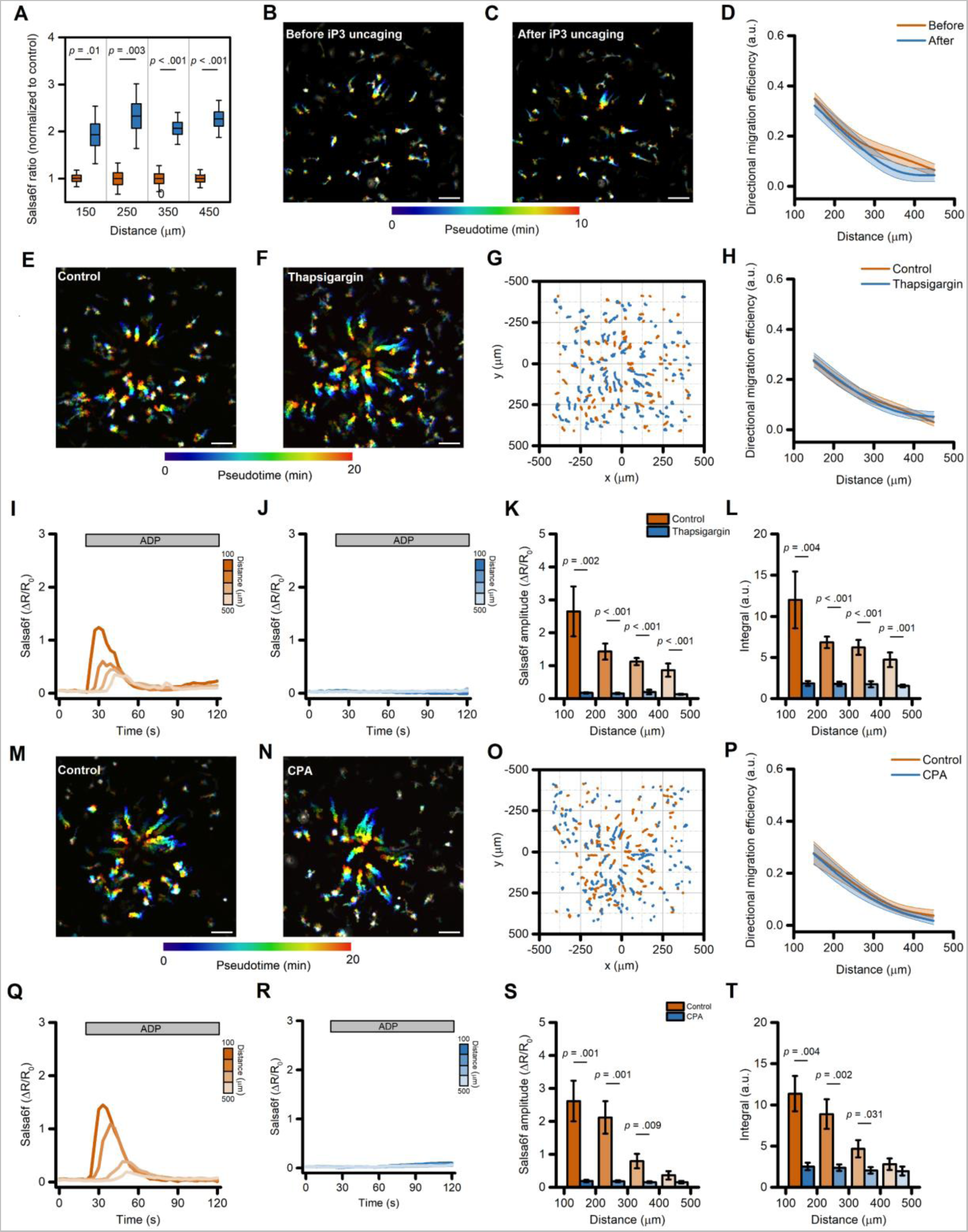
Ca^2+^ release from the ER is not required for directed migration in iMG. (A) Box plots depict quantification of Ca^2+^_i_ levels before (orange) and after (blue) ci-IP_3_ uncaging, averaged at 100 µm radial increments, and expressed as peak amplitude (from n = 7 independent experiments). (B-C) Pseudocolored maximum intensity projection photomicrographs of iMG before (B) and after (C) ci-IP_3_ uncaging while challenged with an ADP gradient. (D) The plot depicts directed migration efficiency towards an ADP gradient of iMG cells before and after ci-IP_3_ uncaging and averaged at 100 µm radial increments (from n = 7 independent experiments). (E-F) Pseudocolored maximum intensity projection photomicrographs of control (E) and thapsigargin-treated (1 µM; F) iMG cells expressing the Salsa6f sensor challenged with an ADP gradient. (G) ‘Flower plot’ depicts 10-min migration tracks as analyzed from the iMG cells in E and F; traces from the two populations were normalized to the pipette tip coordinates and overlaid. (H) Plot depicts directed migration efficiency towards an ADP gradient of control and 1 µM thapsigargin-treated iMG cells averaged at 100 µm radial increments (from n = 7 controls and n = 9 thapsigargin independent experiments). (I-J) Time course of ADP-dependent Ca^2+^_i_ rises as assessed with the Salsa6f sensor. Traces represent average responses at 100 µm radial increments following ADP release from the pipette in control (I) or thapsigargin-treated (J) iMG cells (from n = 7 controls and n = 9 thapsigargin independent experiments). (K) Bar graphs depict Ca^2^ ^+^_i_ amplitude values expressed as ΔR/R_0_. (L) Bar graphs depict Ca^2^ ^+^_i_ integrals, measured as the area under each curve obtained in the two populations. (M-N) Pseudocolored maximum intensity projection photomicrographs of control (M) and CPA-treated (50 µM; F) iMG cells expressing the Salsa6f sensor challenged with an ADP gradient. (O) ‘Flower plot’ depicts 10-min migration tracks as analyzed from the iMG cells in M and N; traces from the two populations were normalized to the pipette tip coordinates and overlaid. (P) The plot depicts directed migration efficiency towards an ADP gradient of control and 50 µM CPA-treated iMG cells averaged at 100 µm radial increments (from n = 6 controls and n = 7 CPA independent experiments). (Q-R) Time course of ADP-dependent Ca^2+^_i_ rises as assessed with the Salsa6f sensor. Traces represent average responses at 100 µm radial increments following ADP release from the pipette in control (Q) or CPA-treated (R) iMG cells (from n = 6 controls and n = 7 CPA independent experiments). (S) Bar graphs depict Ca^2^ ^+^_i_ amplitude values expressed as ΔR/R_0_. (T) Bar graphs depict Ca^2+^_i_ integrals, measured as the area under each curve obtained in the two populations. In D, H, and P a b-spline function was applied for curve smoothing. In D, G, H, U, X, and Y, the comparison of mean values was assessed by a two-tailed unpaired Student’s t-test. In L, P, and Q, comparison of mean values was assessed by one-way ANOVA followed by Tukey’s post-hoc test. Scale bars 100 μm.

Overall, these results confirm that Ca^2+^ released from the ER is required for the early Ca^2+^_i_ elevations following an ADP challenge, but these Ca^2+^_i_ changes are dispensable for directed cell migration.

### ADP-evoked directed microglial migration is mediated by changes in intracellular cAMP concentrations

In early experiments to explore the role of Ca^2+^ signaling in directed motility we employed caffeine (10 mM) as an IP_3_ receptor antagonist [39,40]. As expected for this action, caffeine reduced Ca^2+^_i_ changes triggered by the ADP gradient (Fig. 7A-D). However, in contrast to our other experiments indicating that cell migration is largely independent of Ca^2+^_i_ changes, we unexpectedly found that ADP-evoked directed cell migration was almost completely abrogated by caffeine (Fig. 7E-H), along with significant effects on many other motility parameters (Suppl. Fig. 7A-B).

**Figure 7.**
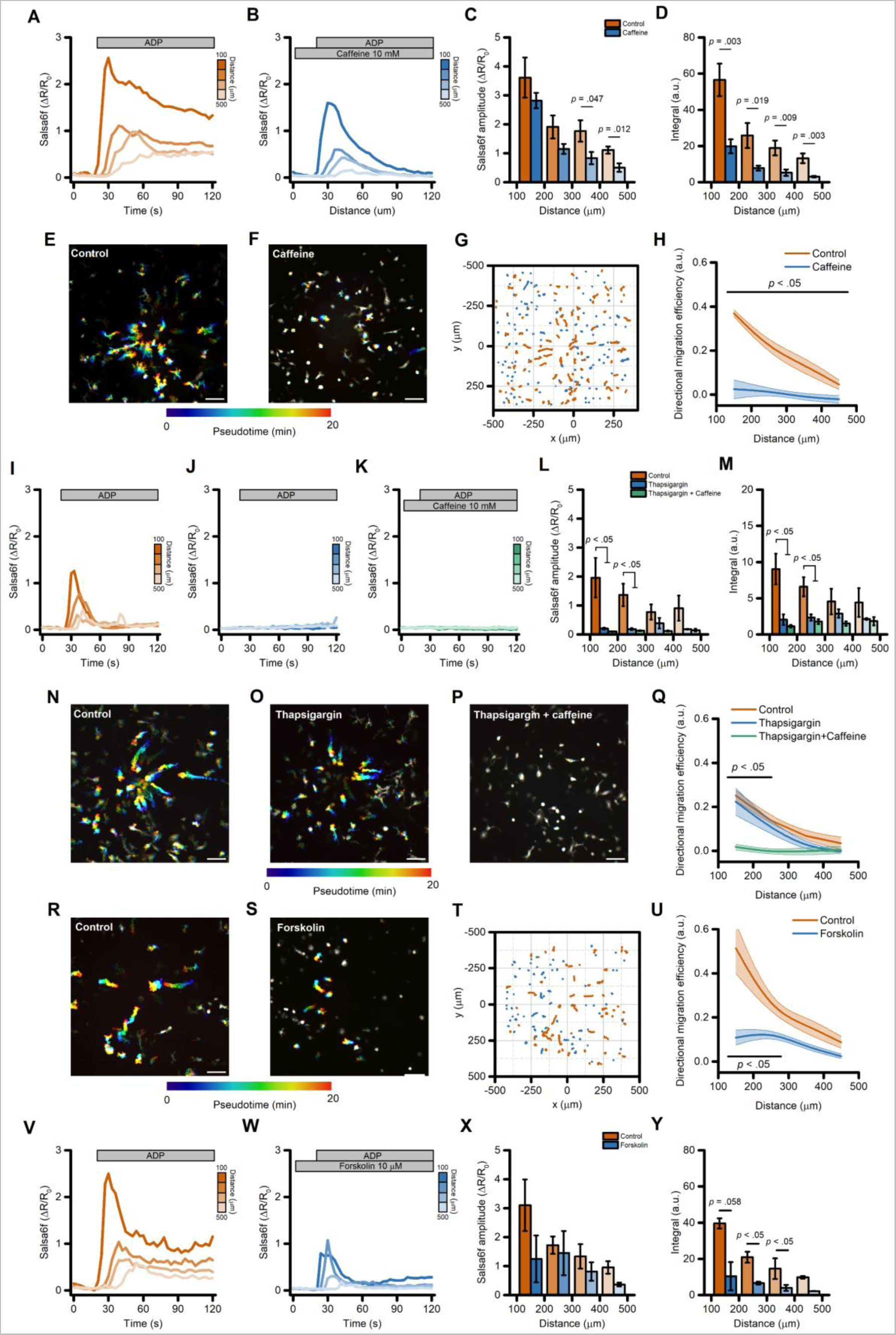
Directed migration of iMG cells is mediated by changes in intracellular cAMP concentrations. (A-B) Pseudocolored maximum intensity projection photomicrographs of control (A) and caffeine-treated (10 mM; B) iMG cells expressing the Salsa6f sensor challenged with an ADP gradient. (C) ‘Flower plot’ depicts 10-min migration tracks as analyzed from the iMG cells in A and B; traces from the two populations were normalized to the pipette tip coordinates and overlaid. (D) The plot depicts directed migration efficiency towards an ADP gradient of control and 10 mM caffeine-treated iMG cells averaged at 100 µm radial increments (from n = 5 controls and n = 6 caffeine independent experiments). (E-F) Time course of ADP-dependent Ca^2+^_i_ rises as assessed with the Salsa6f sensor. Traces represent average responses at 100 µm radial increments following ADP release from the pipette in control (E) or caffeine-treated (F) iMG cells (from n = 5 controls and n = 6 caffeine independent experiments). (G) Bar graphs depict Ca^2+^_i_ amplitude values expressed as ΔR/R_0_. (H) Bar graphs depict Ca^2+^_i_ integrals, measured as the area under each curve obtained in the two populations. (I-K) Pseudocolored maximum intensity projection photomicrographs of control (I), thapsigargin (1 µM; J), and thapsigargin+caffeine-treated (1 µM and 10 mM, respectively; K) iMG cells expressing the Salsa6f sensor challenged with an ADP gradient. (L) Plot depicts directed migration efficiency towards an ADP gradient of the three populations averaged at 100 µm radial increments (from n = 3 controls, n = 3 thapsigargin, and n = 3 thapsigargin+caffeine independent experiments). (M-O) Time course of ADP-dependent Ca^2+^_i_ rises as assessed with the Salsa6f sensor. Traces represent average responses at 100 µm radial increments following ADP release from the pipette in control (M), thapsigargin (N), and thapsigargin+caffeine-treated (O) iMG cells (from n = 3 controls, n = 3 thapsigargin, and n = 3 thapsigargin+caffeine independent experiments). (P) Bar graphs depict Ca^2^ ^+^_i_ amplitude values expressed as ΔR/R_0_. (Q) Bar graphs depict Ca^2^ ^+^_i_ integrals, measured as the area under each curve obtained in the three populations. (R-S) Pseudocolored maximum intensity projection photomicrographs of control (R) and forskolin-treated (10 µM; S) iMG cells expressing the Salsa6f sensor challenged with an ADP gradient. (T) ‘Flower plot’ depicts 10-min migration tracks as analyzed from the iMG cells in R and S; traces from the two populations were normalized to the pipette tip coordinates and overlaid. (U) The plot depicts directed migration efficiency towards an ADP gradient of control and 10 µM forskolin-treated iMG cells averaged at 100 µm radial increments (from n = 2 controls and n = 2 forskolin independent experiments). (V-W) Time course of ADP-dependent Ca^2+^_i_ rises as assessed with the Salsa6f sensor. Traces represent average responses at 100 µm radial increments following ADP release from the pipette in control (V) or forskolin-treated (W) iMG cells (from n = 2 controls and n = 2 forskolin independent experiments). (X) Bar graphs depict Ca^2+^_i_ amplitude values expressed as ΔR/R_0_. (Y) Bar graphs depict Ca^2+^_i_ integrals, measured as the area under each curve obtained in the two populations. In D, L, and U a b-spline function was applied for curve smoothing. The comparison of mean values was assessed by a two-tailed unpaired Student’s t-test. Scale bars 100 μm.

We therefore hypothesized that caffeine affects iMG motility through mechanisms that are independent from Ca^2+^ released from the ER. To test this, we pretreated iMG with thapsigargin and subsequently challenged with caffeine while monitoring ADP-evoked Ca^2+^_i_ changes and directed migration. Results were compared with those obtained from thapsigargin- and sham-treated sister cultures (Fig.7I-Q). In both the thapsigargin- and thapsigargin+caffeine-treated groups, ADP-induced Ca^2+^ changes were abrogated when compared to control cultures (Fig. 7I-M). Notably, only in the presence of caffeine we observed a full impairment of iMG migration properties (Fig. 7P-Q and Suppl. Fig. 7C-D), suggesting that the effect of caffeine on directed motility involves Ca^2+^-independent mechanisms.

At the concentration used (10 mM), caffeine could also act as an inhibitor of phosphodiesterases (PDEs) [41], the enzymes responsible for the degradation of the second messenger cyclic adenosine monophosphate (cAMP). To test whether the intracellular levels of cAMP affect directed cell migration, iMG cells were challenged with the ADP gradient in the presence or absence of forskolin (10 µM), an activator of the cAMP-generating enzyme adenylate cyclase (AC). This approach significantly depressed cell migration and Ca^2+^_i_ changes (Fig. 7R-Y and Suppl. Fig. 7E-F), mirroring observations in caffeine treated cells (Figs 7A-H).

These findings suggest that dysregulation of intracellular levels of cAMP impairs ADP-evoked directed migration in human microglia.

## Discussion

This study has three major findings that provide important insights into the physiology of human microglia. First, exposure to an ADP gradient promotes the directed migration of human microglia, a process abrogated by the pharmacological inhibition of the P2Y_12_ and P2Y_13_ purinergic receptors. Second, chemotaxis of microglia is only marginally affected by pharmacological or genetic blockade of Ca^2+^ signaling pathways acting downstream of ADP-driven P2Y_12_ and P2Y_13_ receptor activation. Third, microglial chemotaxis is significantly impaired upon perturbation of cAMP signaling.

Directed cell migration is a highly complex process that can broadly be summarized into four major steps: the generation of a signal, the ability of a cell to sense that signal, its transduction to the intracellular actors that execute the movement, and the application of oriented forces on extracellular substrates [42]. In this context, our results support the central role played by purinergic metabolites that, when released by damaged cells, act as “find me” chemotactic signals for microglia [43,44]. ADP stimulation profoundly affects the migration properties of iPSC-derived microglia (iMG) cells. Our results are in line with previous reports that identified ADP as a master regulator of microglia motility both *in vitro* and *in vivo* [8].

Chemotactic cues, like those mediated by an ADP gradient, are primarily sensed by ligand-receptor mechanisms [42,45]. Consistent with previous observations in mouse microglia and other cellular systems [4,8,46–50], we recently reported that, in iMG cells, P2Y_12_ and P2Y_13_ act as the primary receptors responsible for sensing ADP, and that their activation has profound effects on several aspects of microglia behavior, including Ca^2+^ signaling, branching, surveillance, and motility [7,26]. We now show that these two receptors are also critical for directed cell migration, extending the range of functions that P2Y_12_ and P2Y_13_ receptors mediate in human microglia. Of note, the two receptors share major commonalities, including similarly high expression levels in microglia, and comparable molecular structure [13,51]. Despite overlapping features, recent studies have started to unravel their distinctive contribution to microglia functioning [13,52]. We did not test the specific impact of each receptor on the migration process because specific agonists/antagonists that can discriminate the contribution of P2Y_12_ and P2Y_13_ receptors are lacking [53]. Genetic manipulation will be informative of the specific contribution provided by each of these purinergic receptors on directed microglia migration.

We also found that iMG cells are sensitive to shear forces and produce Ca^2+^ signals in response to the release of the solution from the gradient-generating pipette with no apparent effect on motility. Preliminary data from our group indicated that these Ca^2+^ changes are extracellular Ca^2+^-dependent [7]. These observations may be explained by the presence of mechano-sensitive, Ca^2+^-permeable PIEZO1 channels in microglia [54,55], whose role in the physiology of these cells has only recently started to be unraveled [54,55].

Mirroring the observations from other cell types – like platelets – the P2Y-mediated effects of ADP were primarily linked to their coupling to G_i_ proteins which, by inhibiting AC, result in decreased cAMP levels [50,56,57]. Recent findings, however, report a more complex scenario in which the receptors are involved in more versatile signaling transduction mechanisms than previously acknowledged. In particular, activation of P2Y_12_ and P2Y_13_ receptors has also been implicated in the PLC/iP_3_/Ca^2+^ pathway, as commonly observed in microglia and consistent with our previous observations [7,58–62]. Unexpectedly, our manipulations to alter intracellular Ca^2+^ signaling had little effect on directed iMG cell migration. Specifically, we found that abrogation of the early Ca^2+^_i_ rise originating from the ER Ca^2+^release or blockade of the sustained, SOCE-dependent Ca^2+^ influx is not necessary for the directed migration of iMG. Consistent with these observations, our results also argue against the contribution of a polarized Ca^2+^_i_ gradient or subtle Ca^2+^_i_ microdomains to directed migration [63,64]. Indeed, the presence of BAPTA (a “shuttle buffer” that promotes the globalization of Ca^2+^ signals [37]) or iP_3_ uncaging to disrupt Ca^2+^_i_ gradients did not appreciably impair migration along an [ADP] gradient. Similarly, experiments performed in iMG cells loaded with EGTA, a slow buffer that isolates local Ca^2+^_i_ signals [37], had no significant effects.

Our findings do not align with several reports indicating a critical role exerted by Ca^2+^_i_ changes in modulating the migration of different cell types, including microglia from rodents [7,34,62–67]. These differences may have several possible explanations. First, the discrepancies may arise from cell type-specific migration mechanisms, as microglia exhibit unique characteristics and functional properties that contribute to distinct migratory behaviors compared to other cell types commonly surveyed in motility studies, like mesenchymal, epithelial, or tumor cells [68]. Second, species-specific differences between human- and rodent model-derived microglia could contribute to variations in migratory responses, thereby emphasizing the importance of studying human-derived cells to interrogate human cell physiology more accurately. This hypothesis is also supported by comparison with previous studies reporting the speed of rodent-derived microglia that were several-fold lower compared to iMG cells in similar settings [33]. A third possible explanation is specific to our experimental setting, as most migration studies are based on suboptimal assays that could mask the specific mechanisms underlying directed migration [33], or that investigate processes more linked to non-directional motility (i.e., random motion following drug application to the bathing medium) [7,68]. In agreement, widely employed chemotaxis assays exhibit deficiencies restricting their applicability in studying cell migration [33]. In particular, the commonly used Boyden (Transwell) chamber and scratch assays lack a stable and controllable chemical gradient, posing limitations for rapidly migrating cells or live imaging experiments, respectively [33]. In this context, the micropipette-based assay employed in this study mitigates these constraints and offers a more flexible and physiologically relevant approach to interrogating cellular dynamics.

Our findings, however, do not completely exclude a contribution of Ca^2+^ in the regulation of directed migration. Remodeling of large processes, protrusion of lamellipodia and filopodia, and ruffling of the leading edge, require Ca^2+^ for effective actin cytoskeleton assembly and elongation [69,70]. The amount of free Ca^2+^ ions available – even in the presence of Ca^2+^ buffers – could be sufficient for these processes.

To promote movement, directional mechanical forces are transferred to the extracellular substrate primarily via Ca^2+^-coordinated integrin-based focal adhesion contact sites [42]. Our results showed that extracellular Ca^2+^ removal had only a modest impact on iMG cell migration properties. A potential explanation for this effect relates to the specific fibronectin-based matrix substrate employed in our model. Early studies demonstrated that cell migration on fibronectin-coated substrates is magnesium (Mg^2+^)-dependent with a supporting role of Ca^2+^ ions [71,72], suggesting that Mg^2+^ ions (present in our Ca^2+^-free medium) can fulfill some of the Ca^2+^-mediated anchoring properties. Besides the dispensable role of ORAI1, our findings of a modest deficit in directed migration in a Ca^2+^-free medium would be compatible with a marginal role played by influx of Ca^2+^ through plasma-membrane TRPV4 and PIEZO1 Ca^2+^ channels expressed by microglia [54,55,73], that have been suggested to be implicated in more specialized motility functions [54,55,73]. In addition, it is unclear whether the modest effect associated with extracellular Ca^2+^ removal is associated with the acute nature of our experimental paradigm, and whether chronic Ca^2+^ withdrawal may result in different outcomes.

To identify ADP-mediated downstream mechanisms involved in iMG migration, we examined the role of cAMP. Pharmacological manipulations with caffeine or forskolin revealed a strong contribution of cAMP signaling in directed cell migration. These observations, together with previously reported findings, suggest cAMP polarization plays a prominent role in controlling microglia dynamics, like surveillance, motility, and directed migration [6,74–76]. Importantly, our findings show that dysregulation of cAMP levels impacts iMG chemotaxis in a Ca^2+^-independent manner.

Besides those described above, another study limitation includes the heterogeneous transcriptional profiles that iMG cells can display *in vitro* [28]. Different transcriptional signatures can reflect diverse microglial states in response to physiologically relevant stimuli that can further influence cell behavior. However, how different microglial states impact the mechanisms regulating directed migration is still largely unexplored.

Taken together, this study provides compelling evidence that Ca^2+^_i_ changes evoked by exposure to purinergic chemoattractants – like ADP – are not necessary for directed microglia migration. These findings raise interesting questions on the significance of Ca^2+^ elevations in the context of microglia functioning. For instance, early Ca^2+^ surges might serve as a kick-off signal for the activation of signaling mechanisms or gene expression changes required by microglia once the injured area is reached. Alternatively, Ca^2+^ rises might support microglia metabolism during lengthy, high energy demanding tasks, like prolonged directed migration or the clearance of cellular debris upon reaching an injury, phenomena that could potentially be missed in the short timeframe of our experimental setting. These findings also highlight the need for further studies to examine the impact of disease-related Ca^2+^ dysregulation and cAMP signaling on microglia functioning.

## Data availability

All the raw images and data from this study are available from the corresponding author upon request.

## Acknowledgments

A.G. is supported by the European Union’s Horizon 2020 research and innovation program under the Marie Sklodowska-Curie grant agreement iMIND – no. 841665 and by the Italian Ministry of University and Research (MUR; PRIN PNRR 2022 – P2022WPRKA). I.P. is supported by the National Institutes of Health (NIH) GM R37 048071. M.B.J. is supported by NIH AG069701 and AG073787. J.P.C. is supported by NIH T32 AG073088. The UCI-ADRC is funded by NIH P30 AG066519. The authors also thank Ram Stanciauskas and Ruby De Casas for their technical assistance.

This work is dedicated to Angelo Veronese.

---

**Supplementary figure 1.**
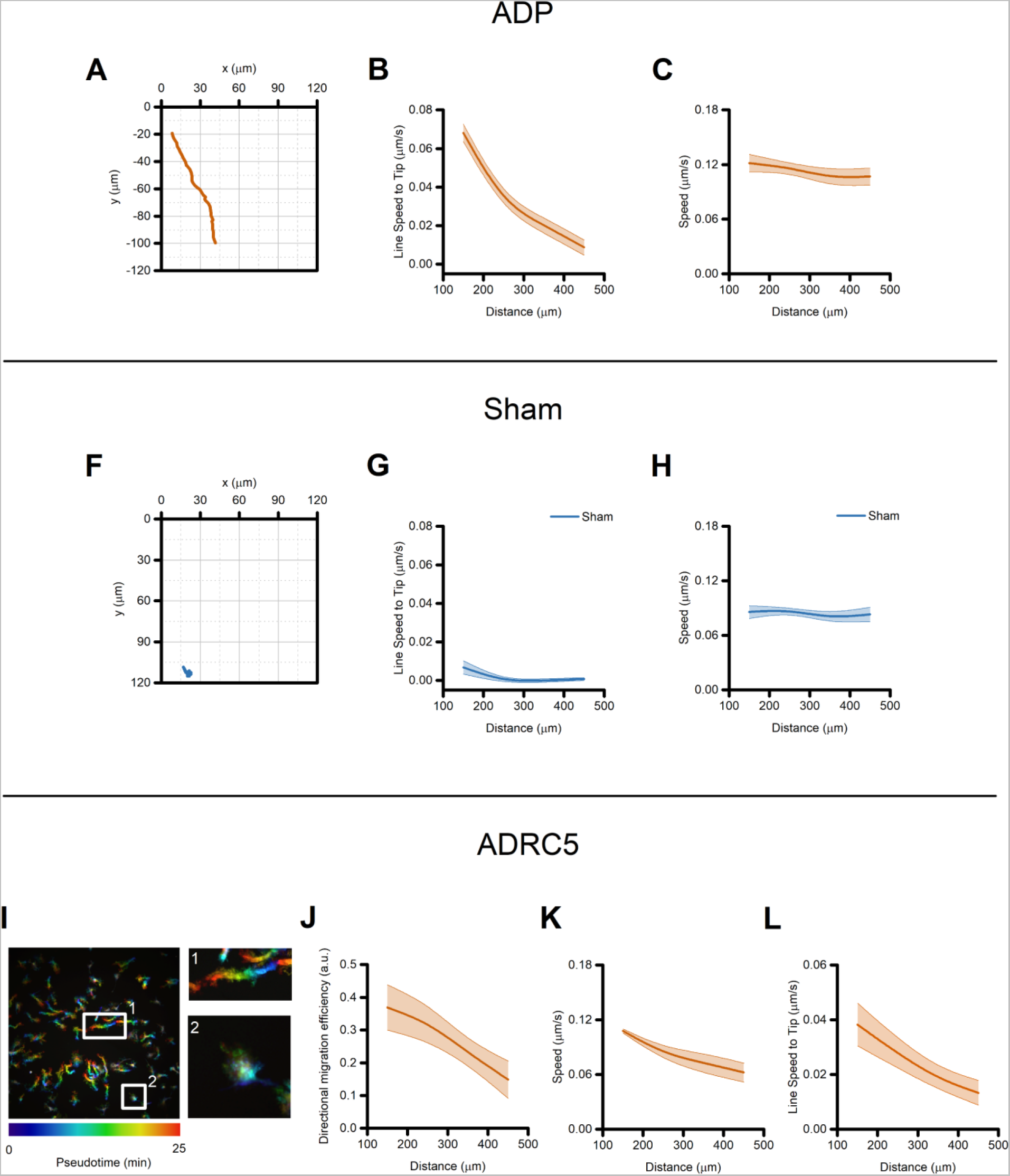
Directed migration of iMG cells following the generation of an ADP gradient. (A) The plot illustrates the trajectory of a representative iMG cell exposed to an ADP gradient whose maximal concentration localizes with the x = 0, y = 0 coordinates. (B) The plot depicts the line speed towards an ADP gradient of iMG cells averaged at 100 µm radial increments (n = 4 independent experiments). (C) The plot depicts speed of ADP-treated iMG cells averaged at 100 µm radial increments (n = 4 independent experiments). Note that speed is almost constant and independent of cell distance from the gradient-generating solution. (F) The plot illustrates the trajectory of a representative iMG cell challenged as in (A) but with ADP being omitted from the gradient-generating solution. (G) The plot depicts the line speed towards the sham gradient of iMG cells averaged at 100 µm radial increments (n = 4 independent experiments). (H) The plot depicts speed of sham-treated iMG cells averaged at 100 µm radial increments (n = 4 independent experiments). (I) Pseudocolored maximum intensity projection photomicrographs of CellTracker Green CMFDA-loaded iMG differentiated from the ADRC5 iPSC line (ADRC5 iMG) and challenged with an ADP gradient. Inlet 1 shows the pseudocolored trajectory of a single ADRC5 iMG cell located in proximity to the gradient. Inlet 2 shows the pseudocolored trajectory of a single ADRC5 iMG cell located distant from the gradient. Note the almost straight pattern followed by the representative cell in Inlet 1 compared to the representative cell in Inlet 2. (J) The plot depicts directed migration efficiency towards an ADP gradient of ADRC5 iMG cells averaged at 100 µm radial increments (n = 4 independent experiments). (K) The plot depicts speed of ADP-treated ADRC5 iMG cells averaged at 100 µm radial increments (n = 4 independent experiments). (L) The plot depicts the line speed towards an ADP gradient of ADRC5 iMG cells averaged at 100 µm radial increments (n = 4 independent experiments).

**Supplementary figure 2.**
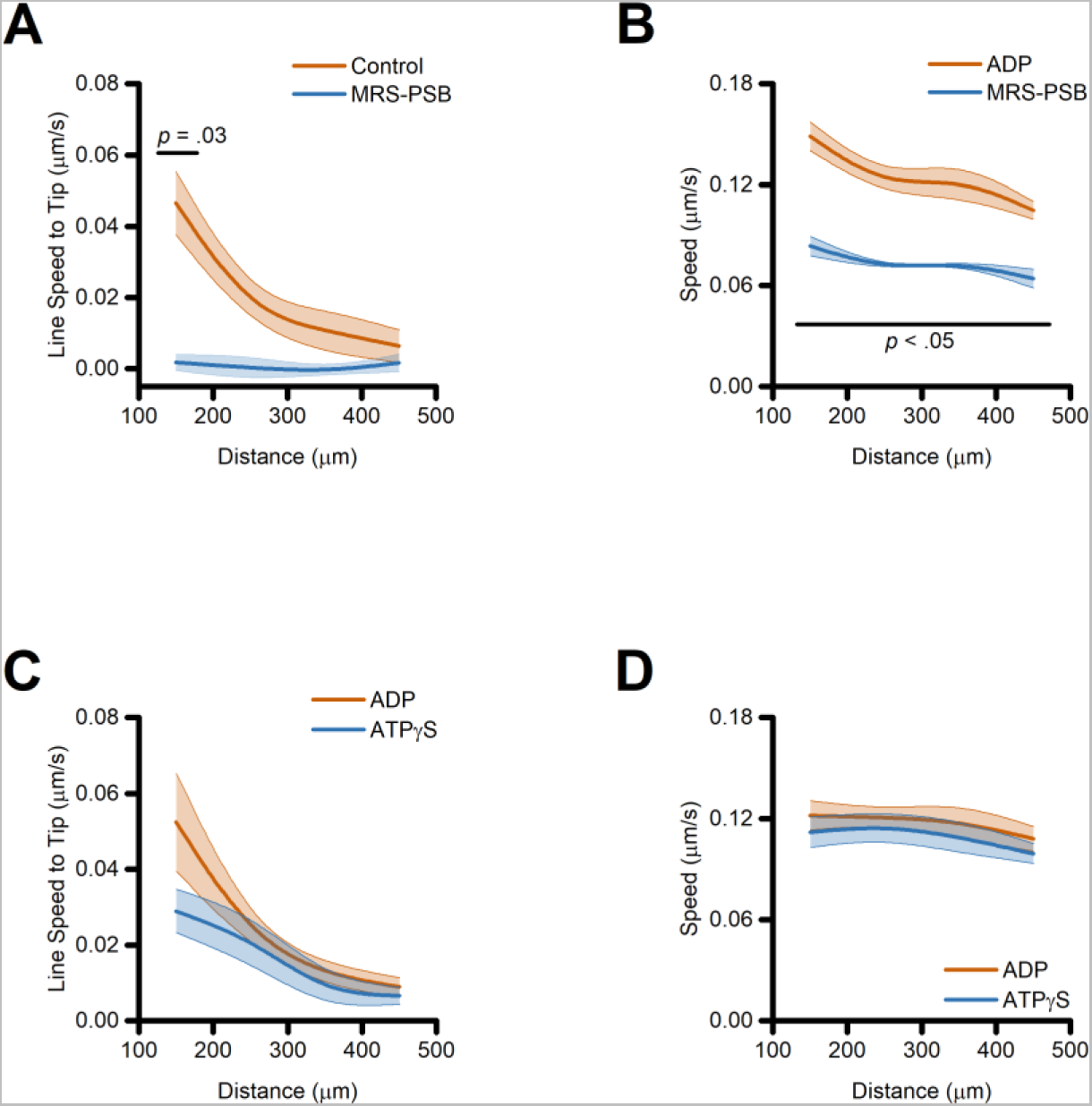
Directed migration of iMG cells is driven by activation of purinergic signaling. (A) The plot depicts line speed towards an ADP gradient of control and PSB 0739 + MRS 2211-treated iMG cells averaged at 100 µm radial increments (from n = 3 control and n = 2 PSB 0739 + MRS 2211 independent experiments). (B) The plot depicts speed of control and PSB 0739 + MRS 2211-treated iMG cells, challenged with ADP, and averaged at 100 µm radial increments (from n = 3 control and n = 2 PSB 0739 + MRS 2211 independent experiments). (C) The plot depicts line speed of iMG cells towards an equimolar ADP or ATPγS gradient and averaged at 100 µm radial increments (from n = 5 ADP and n = 4 ATPγS independent experiments). (D) The plot depicts speed of ADP and ATPγS-challenged iMG cells averaged at 100 µm radial increments (from n = 5 ADP and n = 4 ATPγS independent experiments). In A to D, a b-spline function was applied for curve smoothing. The comparison of mean values was assessed by a two-tailed unpaired Student’s t-test.

**Supplementary figure 3.**
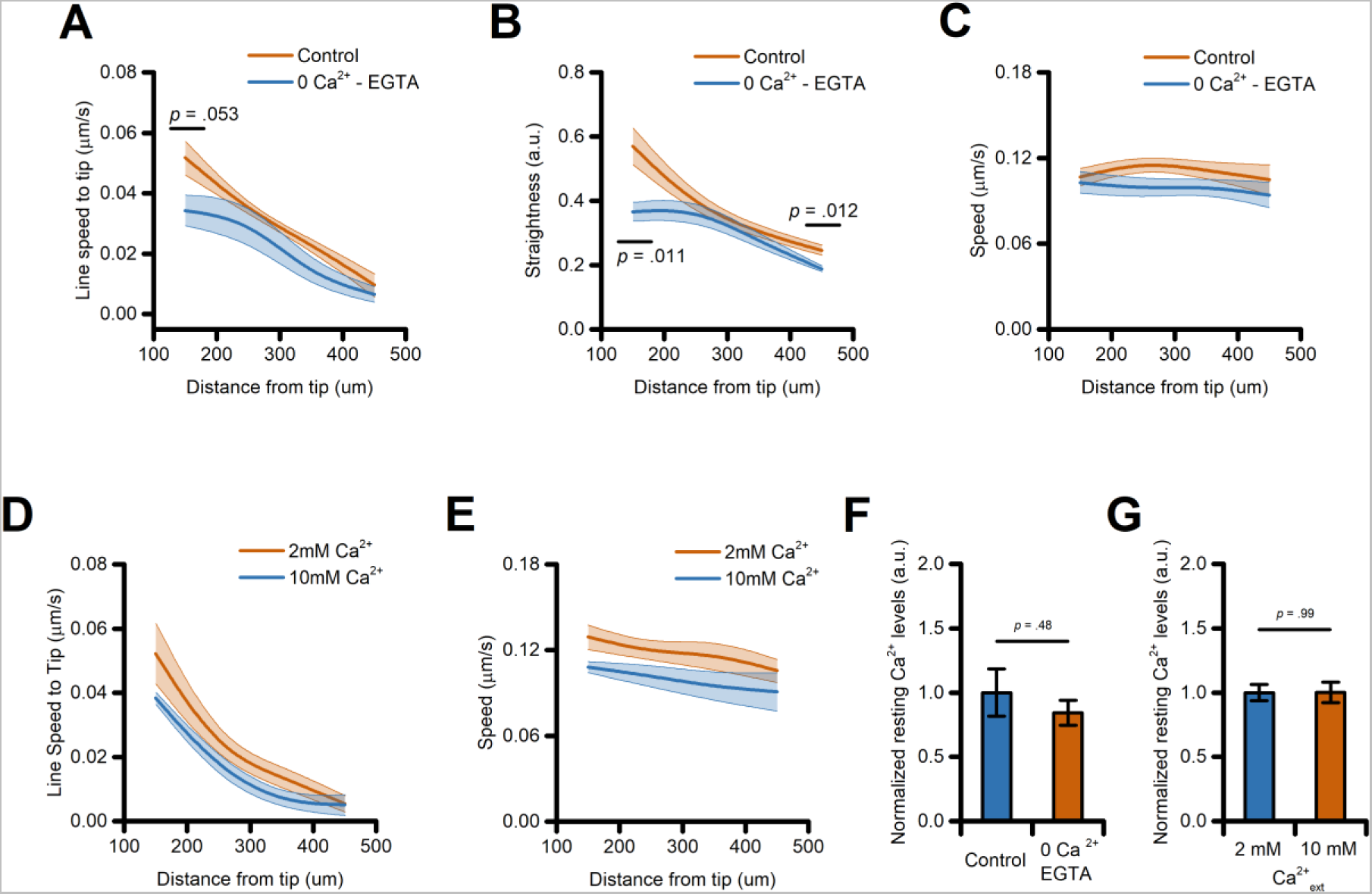
Directed migration of iMG cells is modestly affected by perturbation of the extracellular Ca^2+^ milieu. (A) The plot depicts directed migration efficiency towards an ADP gradient of iMG cells bathed in a control or Ca^2+^-free medium and averaged at 100 µm radial increments (from n = 4 controls and n = 5 Ca^2+^-free independent experiments). (B) The plot depicts straightness of iMG cells bathed in a control or Ca^2+^-free medium, challenged with ADP, and averaged at 100 µm radial increments (from n = 4 controls and n = 5 Ca^2+^-free independent experiments). (C) The plot depicts the speed of iMG cells bathed in a control or Ca^2+^-free medium, challenged with ADP, and averaged at 100 µm radial increments (from n = 4 controls and n = 5 Ca^2+^-free independent experiments). (D) The plot depicts straightness of iMG cells bathed in a control (2 mM Ca^2+^) or 10 mM Ca^2+^-containing medium, challenged with ADP, and averaged at 100 µm radial increments (from n = 5 controls and n = 5 10 mM Ca^2+^ independent experiments). (E) The plot depicts the speed of iMG cells bathed in a control or 10 mM Ca^2+^-containing medium, challenged with ADP, and averaged at 100 µm radial increments (from n = 5 controls and n = 5 10 mM Ca^2+^ independent experiments). (F-G) Bar graphs depict normalized Salsa6f basal ratio values for iMG exposed to either a Ca^2+^-free (F) or a 10mM Ca^2+^-containing (G) extracellular medium (n = 5 independent experiment per condition). Note that the two maneuvers do not affect Ca^2+^_i_ levels. In A to E, a b-spline function was applied for curve smoothing. The comparison of mean values was assessed by a two-tailed unpaired Student’s t-test.

**Supplementary figure 4.**
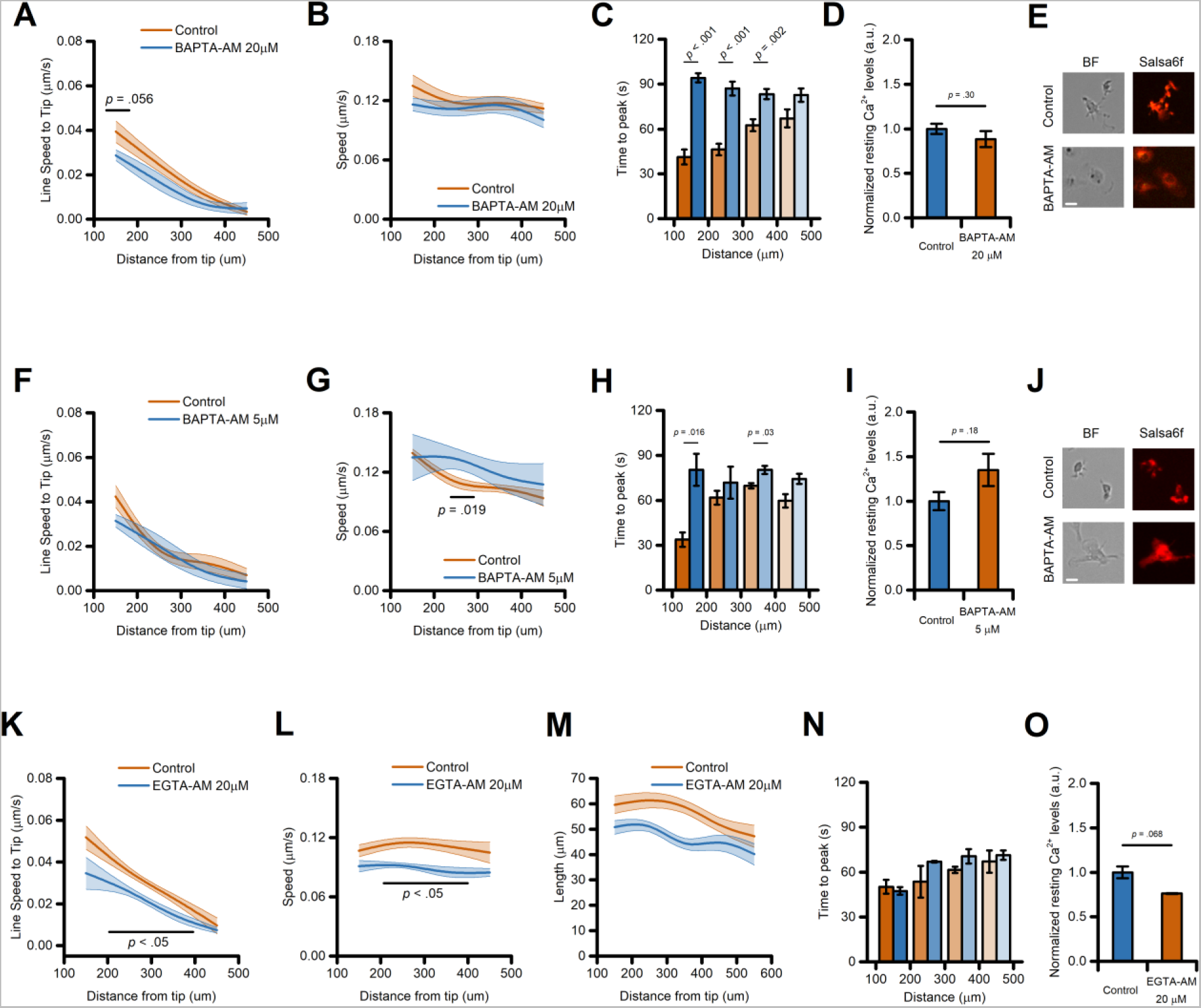
Directed migration of iMG cells is modestly affected by intracellular Ca^2+^ chelators. (A) The plot depicts line speed towards an ADP gradient of control and 20 µM BAPTA-loaded iMG cells averaged at 100 µm radial increments (from n = 5 control and n = 7 20 µM BAPTA independent experiments). (B) The plot depicts speed of control and 20 µM BAPTA-loaded iMG cells, challenged with ADP, and averaged at 100 µm radial increments (from n = 5 control and n = 7 20 µM BAPTA independent experiments). (C) Bar graph depicts ADP-evoked Ca^2+^_i_ kinetics expressed as the time taken by a cell Ca^2+^_i_ concentration to reach its maximum level (time to peak; from n = 5 control and n = 7 BAPTA 20 µM independent experiments). (D) Bar graphs depict normalized Salsa6f basal ratio values for control and 20 µM BAPTA-loaded iMG (from n = 5 control and n = 7 20 µM BAPTA independent experiments). (E) Representative brightfield (BF) and fluorescent (Salsa6f) photomicrographs of control and 20 µM BAPTA-loaded iMG cells. Note the flattened, less branched morphology displayed by BAPTA-treated iMG cells at the end of the loading procedure. (F) The plot depicts line speed towards an ADP gradient of control and 5 µM BAPTA-loaded iMG cells averaged at 100 µm radial increments (from n = 3 control and n = 3 5 µM BAPTA independent experiments). (G) The plot depicts speed of control and 5 µM BAPTA-loaded iMG cells, challenged with ADP, and averaged at 100 µm radial increments (from n = 3 control and n = 3 5 µM BAPTA independent experiments). (H) Bar graph depicts ADP-evoked Ca^2+^_i_ kinetics expressed as the time taken by a cell Ca^2+^_i_ concentration to reach its maximum level (time to peak; from n = 3 control and n = 3 5 µM BAPTA independent experiments). (I) Bar graphs depict normalized Salsa6f basal ratio values for control and 5 µM BAPTA-loaded iMG (from n = 3 control and n = 3 5 µM BAPTA independent experiments). (J) Representative brightfield (BF) and fluorescent (Salsa6f) photomicrographs of control and 5 µM BAPTA-loaded iMG cells. (K) The plot depicts line speed towards an ADP gradient of control and 20 µM EGTA-loaded iMG cells averaged at 100 µm radial increments (from n = 4 control and n = 4 20 µM BAPTA independent experiments). (L) The plot depicts speed of control and 20 µM EGTA-loaded iMG cells, challenged with ADP, and averaged at 100 µm radial increments (from n = 4 control and n = 4 20 µM EGTA independent experiments). (M) The plot depicts the total length travelled by control and 20 µM EGTA-loaded iMG cells, challenged with ADP, and averaged at 100 µm radial increments (from n = 3 control and n = 3 20 µM EGTA independent experiments). (N) Bar graph depicts ADP-evoked Ca^2+^_I_ kinetics expressed as the time taken by a cell Ca^2+^_I_ concentration to reach its maximum level (time to peak; from n = 3 control and n = 3 20 µM EGTA independent experiments). (O) Bar graphs depict normalized Salsa6f basal ratio values for control and 20 µM BAPTA-loaded iMG (from n = 3 control and n = 3 20 µM EGTA independent experiments). In A, B, F, G, and K-M a b-spline function was applied for curve smoothing. The comparison of mean values was assessed by a two-tailed unpaired Student’s t-test. Scale bars 20 μm.

**Supplementary figure 5.**
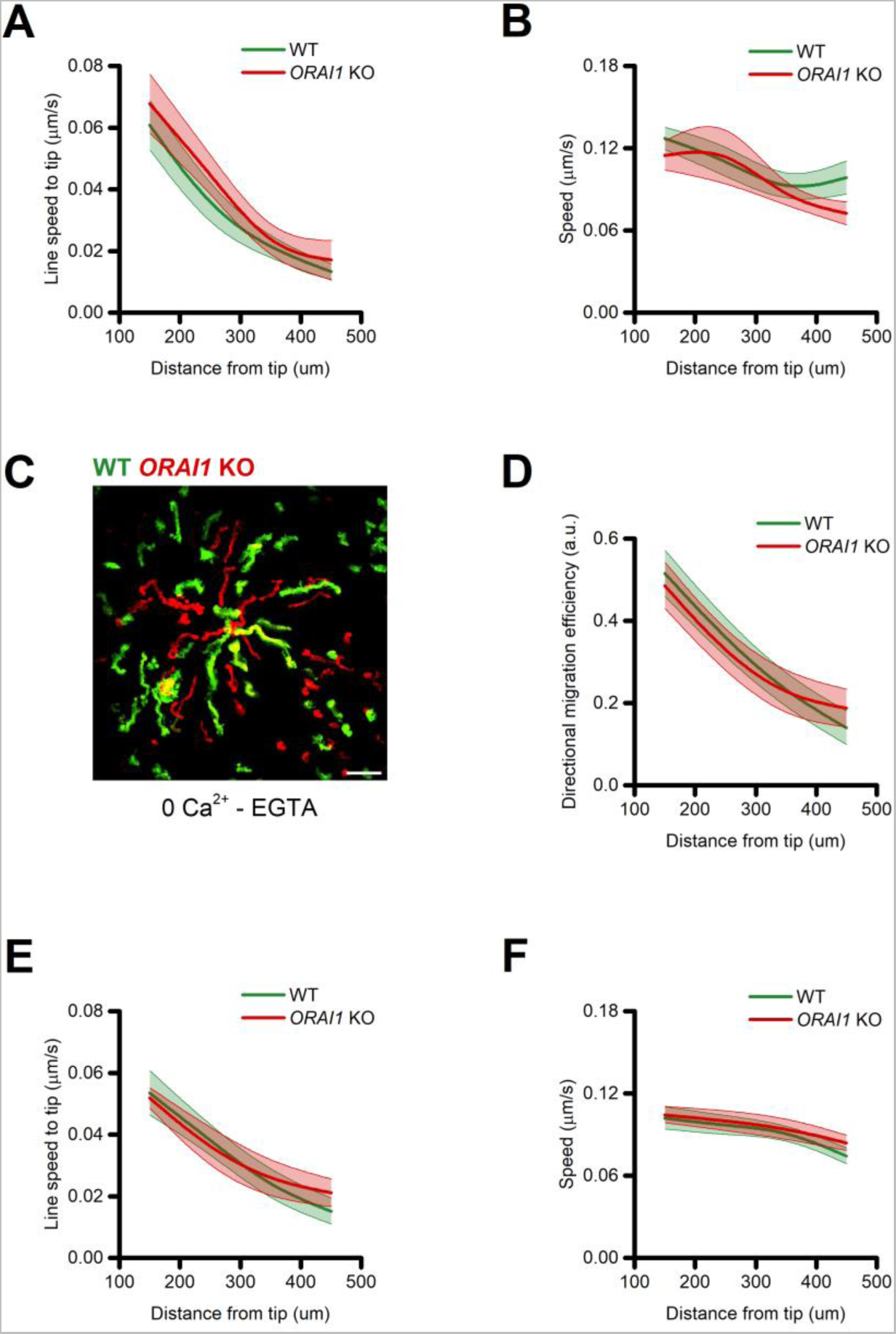
SOCE is not required for directed migration of iMG cells. (A) The plot depicts line speed towards an ADP gradient of WT and *ORAI1* KO iMG cells averaged at 100 µm radial increments (from n = 6 independent experiments). (B) The plot depicts speed of WT and *ORAI1* KO iMG cells, challenged with ADP, and averaged at 100 µm radial increments (from n = 6 independent experiments). (C) Maximum intensity projection photomicrographs of WT (Green) and *ORAI1* KO (Red) iMG cells challenged with an ADP gradient in Ca^2+^-free medium. (D) The plot depicts directed migration efficiency towards an ADP gradient in Ca^2+^-free medium of WT and *ORAI1* KO cells averaged at 100 µm radial increments (from n = 7 independent experiments). (E) The plot depicts line speed towards an ADP gradient in Ca^2+^-free medium of WT and *ORAI1* KO cells averaged at 100 µm radial increments (from n = 7 independent experiments). (F) The plot depicts speed of WT and *ORAI1* KO iMG cells, challenged with ADP in a Ca^2+^-free medium, and averaged at 100 µm radial increments (from n = 7 independent experiments). In A, B, and D-F a b-spline function was applied for curve smoothing. The comparison of mean values was assessed by a two-tailed unpaired Student’s t-test. Scale bar 100 μm.

**Supplementary figure 6.**
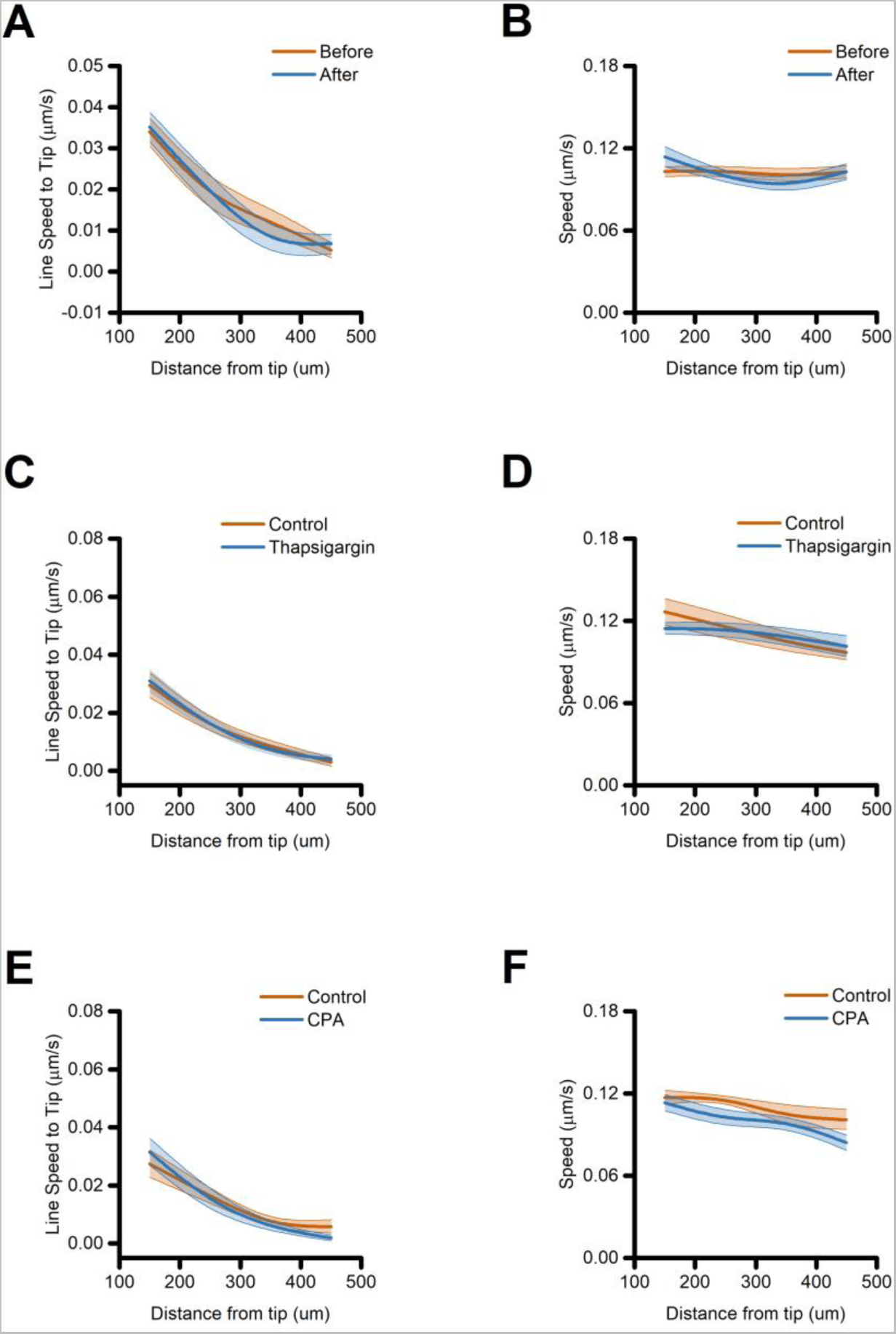
Ca^2+^ release from the ER is not required for directed migration in iMG. (A) The plot depicts line speed towards an ADP gradient of iMG cells before and after ci-IP_3_ uncaging and averaged at 100 µm radial increments (from n = 7 independent experiments). (B) The plot depicts speed of iMG cells before and after ci-IP_3_ uncaging, challenged with ADP, and averaged at 100 µm radial increments (from n = 7 independent experiments). (C) The plot depicts line speed towards an ADP gradient of control and 1 µM thapsigargin-treated iMG cells averaged at 100 µm radial increments (from n = 7 controls and n = 9 thapsigargin independent experiments). (D) The plot depicts speed of control and 1 µM thapsigargin-treated iMG cells, challenged with ADP, and averaged at 100 µm radial increments (from n = 7 controls and n = 9 thapsigargin independent experiments). (E) The plot depicts line speed towards an ADP gradient of control and 50 µM CPA-treated iMG cells averaged at 100 µm radial increments (from n = 6 controls and n = 7 CPA independent experiments). (F) The plot depicts speed of control and 50 µM CPA-treated iMG cells, challenged with ADP, and averaged at 100 µm radial increments (from n = 6 controls and n = 7 CPA independent experiments). A b-spline function was applied to all plots for curve smoothing. The comparison of mean values was assessed by a two-tailed unpaired Student’s t-test.

**Supplementary figure 7.**
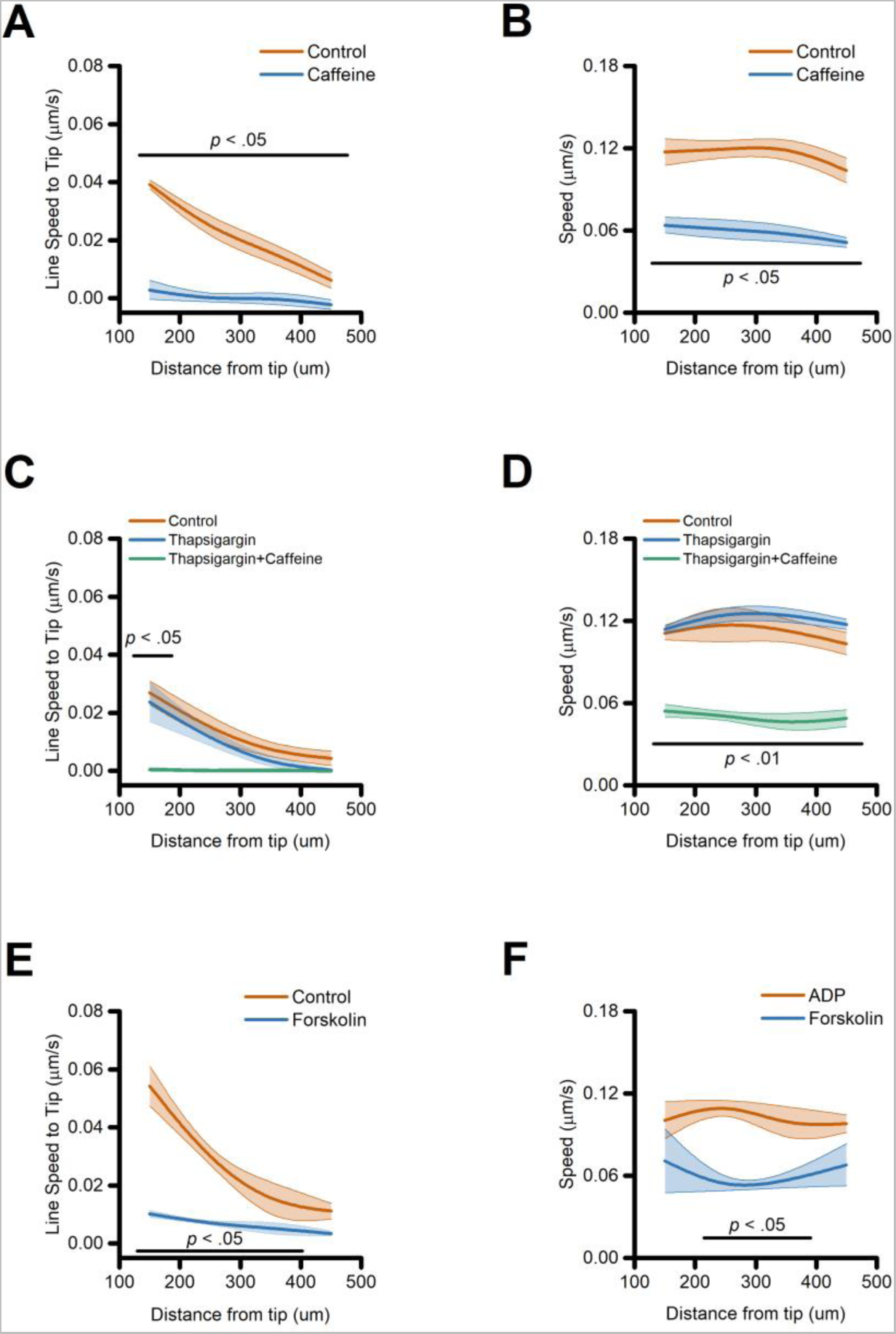
Directed migration of iMG cells is mediated by changes in intracellular cAMP concentrations. (A) The plot depicts line speed towards an ADP gradient of control and 10 mM caffeine-treated iMG cells averaged at 100 µm radial increments (from n = 5 controls and n = 6 caffeine independent experiments). (B) The plot depicts speed of control and 10 mM caffeine-treated iMG cells, challenged with ADP, and averaged at 100 µm radial increments (from n = 5 controls and n = 6 caffeine independent experiments). (C) The plot depicts line speed towards an ADP gradient of control, thapsigargin, and thapisgargin+caffeine-treated iMG cells averaged at 100 µm radial increments (from n = 3 controls, n = 3 thapsigargin, and n = 3 thapsigargin+caffeine independent experiments). (D) The plot depicts speed of the three populations, challenged with ADP, and averaged at 100 µm radial increments (from n = 3 controls, n = 3 thapsigargin, and n = 3 thapsigargin+caffeine independent experiments). (E) The plot depicts line speed towards an ADP gradient of control and 10 µM forskolin-treated iMG cells averaged at 100 µm radial increments (from n = 2 controls and n = 2 forskolin independent experiments). (F) The plot depicts speed of control and 10 µM forskolin-treated iMG cells, challenged with ADP, and averaged at 100 µm radial increments (from n = 2 controls and n = 2 forskolin independent experiments). A b-spline function was applied to all plots for curve smoothing. The comparison of mean values was assessed by a two-tailed unpaired Student’s t-test.

**Supplementary Movie 1.** The video shows a feasibility experiment demonstrating simultaneous imaging of directed migration and intracellular Ca^2+^_i_ in human microglia. The video features iMG expressing the Salsa6f Ca^2+^ reporter exposed to an ADP gradient. This time series highlights the ability to monitor both Ca^2+^_i_ levels and iMG dynamics over an extended period. Video acquired at 0.3 Hz. Scale bar 100 µm.

**Supplementary Movie 2.** The video shows a feasibility experiment for multiplex imaging of human microglia directed migration. The video exhibits two iMG populations loaded with distinct CellTracker dyes exposed to an ADP gradient. The time series illustrates the simultaneous, long-term tracking of iMG dynamics. Video acquired at 0.03 Hz. Scale bar 100 µm.

## Notes

### Competing Interest Statement

The authors have declared no competing interest.

